# Disease exacerbation by fibroblast inclusion in Duchenne Muscular Dystrophy MYOrganoids reveals limitations of microdystrophin therapeutic efficacy

**DOI:** 10.1101/2023.07.26.550063

**Authors:** Laura Palmieri, Louna Pili, Abbass Jaber, Ai Vu Hong, Matteo Marcello, Riyad El-Khoury, Guy Brochier, Anne Bigot, David Israeli, Isabelle Richard, Sonia Albini

**Affiliations:** Genethon, 91100 Evry, France; Université Paris-Saclay, Univ Evry, Inserm, Généthon, Integrare research unit UMR_S951, 91000, Evry, France; Institut de Myologie, Neuromuscular Morphology Unit, Groupe Hospitalier Pitié-Salpêtrière, Paris, France; AP-HP, Centre de Référence de Pathologie Neuromusculaire Nord/Est/Ile de France, Groupe Hospitalier Pitié-Salpêtrière, Paris, France; Sorbonne Université, Inserm, Institut de Myologie, Centre de Recherche en Myologie, F-75013 Paris, France

**Keywords:** iPSC, DMD, fibroblasts, fibrotic, MYOrganoids, AAV, microdystrophin

## Abstract

Current gene therapy approaches for Duchenne muscular dystrophy (DMD) using AAV-mediated delivery of microdystrophin (µDys) have shown limited efficacy in patients, contrasting with the favorable outcomes observed in animal models. This discrepancy is partly due to the lack of models that replicate key pathogenic features associated with the severity of the human disease, such as fibrosis and muscle dysfunction. To tackle the translational gap, we develop a human disease model that recapitulates these critical hallmarks of DMD for a more predictive therapeutic investigation. Using a muscle engineering approach, we generate MYOrganoids from iPSC-derived muscle cells co-cultured with fibroblasts that enable functional maturation for muscle force analysis upon contractions. Incorporation of DMD fibroblasts within DMD iPSC-derived muscle cells allows phenotypic exacerbation by unraveling of fibrotic signature and fatiguability through cell-contact-dependent communication. Although µDys gene transfer partially restores muscle resistance, it fails to fully restore membrane stability and reduce profibrotic signaling. These findings highlight the persistence of fibrotic activity post-gene therapy in our human DMD system, an unparalleled aspect in existing DMD models, and provide the opportunity to explore the underlying mechanisms of dysregulated cellular communication to identify anti-fibrotic strategies empowering gene therapy efficacy.

## Introduction

Duchenne muscular dystrophy (DMD; ONIM: #310200) is an X-linked disorder that affects one in every 5000 male births ^1^ with no resolutive cure up to date. It is characterized by progressive muscle wasting affecting skeletal muscles primarily and cardiac and respiratory muscles later, thereby causing premature death ^2^. DMD is caused by genetic mutations in the *DMD* gene, leading to the absence of Dystrophin, an essential protein that provides physical support to myofibers by linking them to the extracellular matrix through the Dystrophin Glycoprotein Complex (DGC) ^3–6^. The lack of Dystrophin results in a series of muscle membrane breakdowns and repairs, leading to subsequent secondary issues like chronic inflammation and fibrosis ^7–10^. Fibrosis is an excessive deposition of extracellular matrix components like fibronectin and collagen, triggered by overactivation of Transforming Growth Factor beta (TGF-β) and leading to loss of muscle functionality ^12–13^. Besides being a critical driver of DMD progression, fibrosis also hampers gene therapy efficacy and is therefore paramount to counteract this process that is well-established in patients.

Gene therapy using adeno-associated virus (AAV) is currently the most promising treatment for Duchenne muscular dystrophy. Ongoing clinical trials use AAV to deliver short forms of Dystrophin, known as microdystrophin (µDys), which encodes a truncated but functional protein ^13–19^. However, while the therapeutic effects were unequivocally achieved in DMD animal models ^20^, the results from clinical trials revealed only partial therapeutic efficacy in terms of gain of muscle function and rarely addressed whether fibrotic activity and signaling were reduced by gene transfer ^21,22^. These observations confirm the limited translatability of results obtained in animal models to human patients. It appears therefore crucial to develop time and cost-effective high throughput models, mimicking the severity of human DMD pathology, suitable for research investigation and therapeutic screening.

In this context, *in vitro* modeling based on human cells is a valuable option. In particular, the induced pluripotent stem cells (iPSC) technology offers the opportunity to derive an unlimited number of specialized cells from patients for disease modeling and drug screening ^23,24^. Among the *in vitro* cellular models, organoid-like structures are becoming invaluable for disease modeling as the use of 3D cultures and biomaterials allows the reconstitution of tissue architecture and microenvironment that are instrumental for pathophysiological evaluations ^25,26^. Tissue engineering applications for AAV gene therapy have been exploited mostly in the context of retinopathies ^24,27–29^ while only limitedly explored for muscular disorders. Hence, having human DMD models is of utmost importance to advance gene therapy and provide a platform for predictive screening. Although several *in vitro* 3D systems are accessible for modeling Duchenne muscular dystrophy ^30–33^, their throughput use is limited by the long duration, variability, related to the complexity of cellular composition achieved, and lack of disease-specific readouts for muscle function.

Here, we report on the generation of iPSC-derived muscle organoid structure, named hereafter MYOrganoids. We employ and adapt an engineered muscle platform to generate MYOrganoids using a previously reported method for direct iPSC conversion into 2D skeletal muscle cells ^34–36^. As a strategy to increase the structural and functional maturation required for pathophysiological studies, we use fibroblasts as they are a major source of connective tissue which is a key regulator of differentiation and muscle structure. Moreover, fibroblasts act as a source of microenvironment cues exerted by their secretory activity and they are therefore regulators of the muscle niche that undergoes pathological remodeling during disease ^37,38^ .Here we show that fibroblast inclusion enhances the structural and functional maturation of the muscle cells. In a DMD context, fibroblasts allow exacerbation of phenotypic traits by direct interaction with muscle cells and reveal key hallmarks of DMD such as fibrosis and muscle weakness over repeated contractions.

Our study also evaluates for the first time the therapeutic efficacy of AAV-mediated µDys gene transfer, in engineering muscle tissues, as proof of concept of their suitability for studying disease mechanisms and evaluating potential therapeutics. By using different doses of µDys in DMD MYOrganoids, we observed a dose-dependent response in restoring muscle function while only a partial effect at the level of membrane stability and fibrotic signature in DMD muscles. Our findings indicate that patient-derived MYOrganoids, whose pathogenic traits are exacerbated, are suitable for studying the fibrotic process orchestrated by either muscle or fibroblast population and its interplay with gene transfer approaches. Our system has therefore the potential to identify molecular mechanisms driving the dystrophic process and accelerate the identification of effective therapeutics for DMD.

## Results

### Generation of structurally organized 3D human MYOrganoids by direct conversion of iPSC and inclusion of fibroblasts

MYOrganoids were generated from human iPSC committed to differentiating into the myogenic lineage by inducible expression of MyoD and BAF60C ^34^ (referred to as iPSC^BM.^) that can directly generate myotubes. MYOrganoids were prepared starting from iPSC^BM^ after one day from the induction of myogenic genes. The casting procedure was performed through adaptation of an engineered muscle system ^33,39^ which results in the growth of the tissue in a ring format supported by two flexible silicon stretchers. The 3D cultures were kept for 2 days in growth medium, afterwards medium was replaced for differentiation for another 12 days (**Figure 1A**). The differentiation protocol was optimized from the conditions previously reported ^35,36,40^ using myogenic commercial media that ensured the highest expression of myogenic markers and myogenic differentiation in monolayer conditions (**Figure S1A-B**). Since cellular heterogenicity, especially of mesenchymal origin, is important for muscle formation ^33,37,38,41^, we included human fibroblasts during the casting procedure, to assess whether this would affect muscle organization. For this aim, casting was performed using iPSC^BM^ cells in the presence or absence of human fibroblasts. Achieving alignment and differentiation simultaneously necessitates a delicate balance between fibroblasts and muscle cells (as noted by N. Rao et al., 2013). To replicate physiological conditions accurately, we incorporated a fibroblast concentration that mirrors the stromal population detectable through single cell and single nuclei-RNA seq analysis of muscles ^42–45^. We found that including 10% fibroblasts, accelerated the condensation and growth over time of the muscle rings into a compact structure 0.8 mm long and 1 mm thick at day 14 (**Figure 1B**). By performing immunofluorescence in whole-mount tissues for sarcomeric α-actinin (SAA), we could detect an enrichment of SAA-positive myotubes throughout the ring-shaped micro-tissue (**Figure 1C**). We then aimed to assess the impact of fibroblast inclusion on muscle structure. Since the organization of muscle cells within the ECM plays a key role in fusion and maturation ^46^, both alignment of myotube and circularity, a consequence of their parallelism, were evaluated. Staining for myosin heavy chain (MyHC, myotube marker) and vimentin (fibroblasts marker) on longitudinal sections showed fibroblast recruitment near muscle fibers (**Figure 1D**). Myotubes alignment was determined by measuring the angles in between myotubes ^47^ and showed a significant decrease towards 0 degrees upon fibroblast inclusion, which indicates parallelism while, in the MYOrganoids without fibroblasts, we detected a disordered pattern of muscular cells (**Figure 1E**). Additionally, the circularity of myofibers was measured from transversal sections stained for the membrane marker wheat germ agglutinin (WGA), using the ratio between X and Y Feret diameters (**Figure 1D**). MYOrganoids including fibroblasts had an improved circularity (ratio closer to 1) when compared to control (**Figure 1E**). Improved myotube circularity and alignment as shown, indicate that fibroblast incorporation during the casting procedure guides skeletal cell orientation providing structural support for MYOrganoids, a prerequisite for maturation.

**Figure 1.**
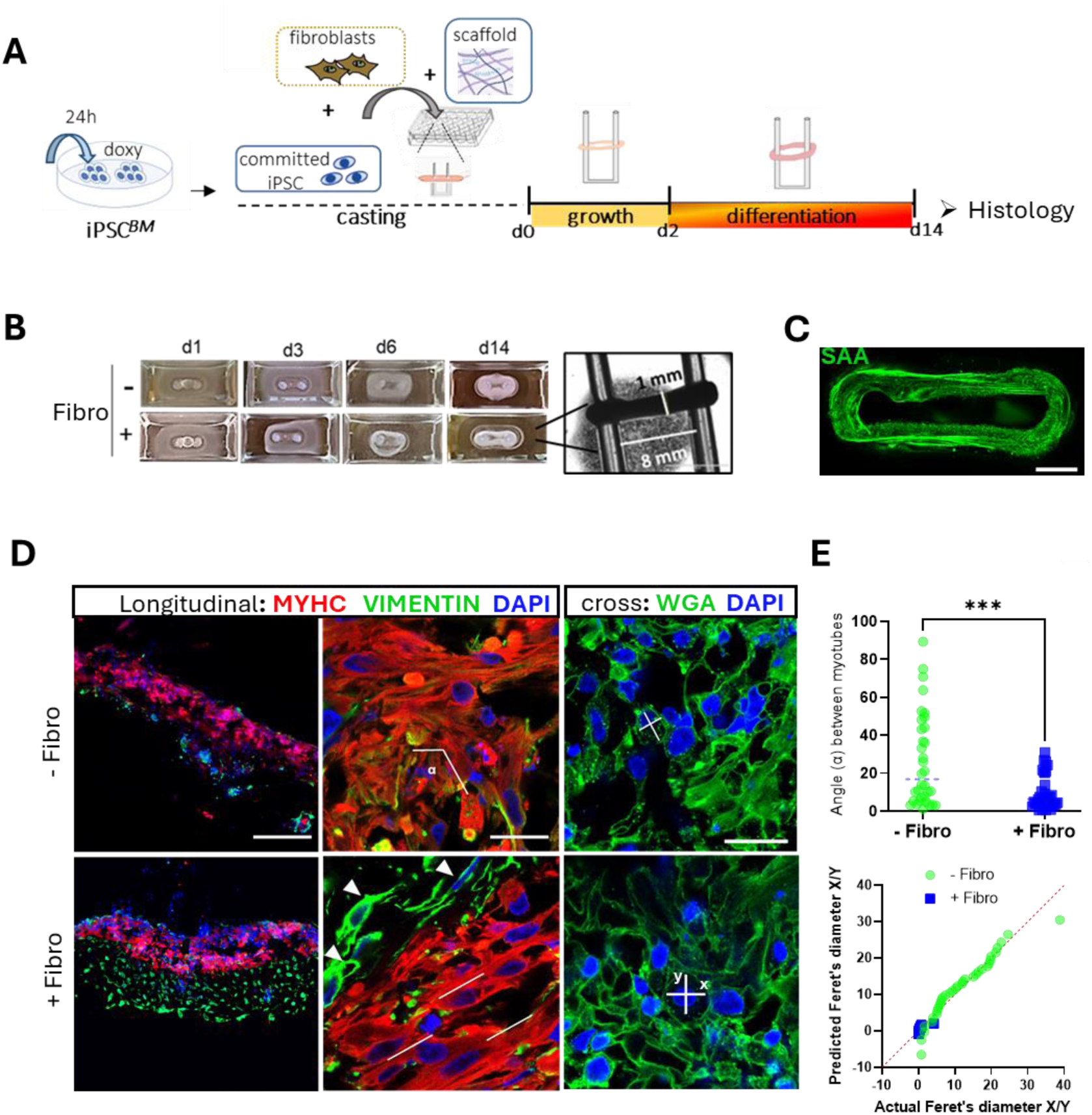
Generation of iPSC-derived MYOrganoids and impact of fibroblast inclusion on muscle organization. **(A)** Scheme of the protocol used to generate muscle artificial tissues (MYOrganoids) from iPSC committed towards the myogenic lineage by 24h treatment with doxycycline for inducible expression of Myod and BAF60C transgenes (iPSC^BM^). The casting procedure included: committed iPSC, fibroblasts when indicated (+/-fibro) and a collagen-based scaffold, within a 48-well plate equipped with silicon pillars. After 2 days in growth medium, the 3D structures were shifted to differentiation medium until day 14 for histological analysis. (**B**) Condensation kinetics of MYOrganoids +/-fibro. (**C**) Whole-mount staining of MYOrganoids with SARCOMERIC α-ACTININ (SAA) and 3D reconstruction of the ring-shaped constructs using confocal imaging. Scale bar: 1 mm. **(D)** Representative longitudinal and cross sections of MYOrganoids +/− fibroblasts, immunostained for Myosin Heavy Chain (MYHC) and VIMENTIN or for wheat germ agglutinin (WGA). Nuclei were visualized with DAPI. Scale bars 200 µm (left panel) and 10 µm (middle and right panel). Arrows indicate fibroblasts recruited adjacently to the muscle fibers; α is the angle formed between myotubes; lines indicate aligned myotubes, crosses represent X/Y myotubes diameters. **(E)** The alignment was calculated based on the angle (α) formed between myotubes (α close to 0 corresponds to aligned myotubes, while far from 0 corresponds to not aligned myotubes), while circularity from X/Y myotubes diameters ratio (ratio 1 circular, far from 1 not circular). Ratio X/Y diameter is represented by a QQ normality plot. Data were collected from 3 independent experiments with n=3.Unpaired two-tailed t-test was used (***p ≤ 0.001).

### Increased structural and functional maturation of fibroblast-including MYOrganoids

Since muscle maturation is strictly dependent on the internal myofiber organization, we evaluated the differentiation of our 3D MYOrganoids by looking at the sarcomere structure. Transversal and longitudinal sections were used to monitor dystrophin (DYS) expression at the sarcolemma (**Figure 2A**) and sarcomeric α-actinin (SAA) localization for assessment of the striation pattern typical of mature myotubes (**Figure 2B**). Dystrophin was properly localized to the muscle membrane of myotubes from MYOrganoids including fibroblasts and was significantly more expressed than MYOrganoids without fibroblasts (**Figure 2D**). Remarkably, the maturation index, reported as a percentage of the number of nuclei included in striated myotubes, was significantly superior in MYOrganoids including fibroblasts as compared to MYOrganoids without fibroblasts which appear very disorganized with a rare appearance of striations (**Figure 2E**). Increased maturation of myotubes within MYOrganoids including fibroblast was also supported by analysis of the fusion index, indicating a significantly higher percentage of multinucleated myotubes (>3 nuclei) and a lower percentage of mononucleated ones (**Figure 2F**). This evidence highlights the positive role of fibroblasts in the maturation process through fusion and multinucleation. The proper sarcomeric organization was also confirmed by electron microscopy where we could detect longer, properly formed Z patterning and the presence of I and A bands along with an overall increase of sarcomeric density and alignment (**Figure 2C**). Consistently, MYOrganoids containing fibroblasts showed wider Z-line (**Figure 2H**), index of higher maturation of sarcomeres ^48^. We further performed gene expression analysis for terminal differentiation markers such as muscle creatine kinase (*MCK*), myosin heavy chain (*MYH*) isoforms, such as *MYH2*, representative of fast adult fiber type, and *MYH7*, as a slow fibers marker (**Figure 2G**). Higher expression of all genes in MYOrganoids containing fibroblasts confirms the acquisition of a more mature state, compared to MYOrganoids without fibroblasts.

**Figure 2.**
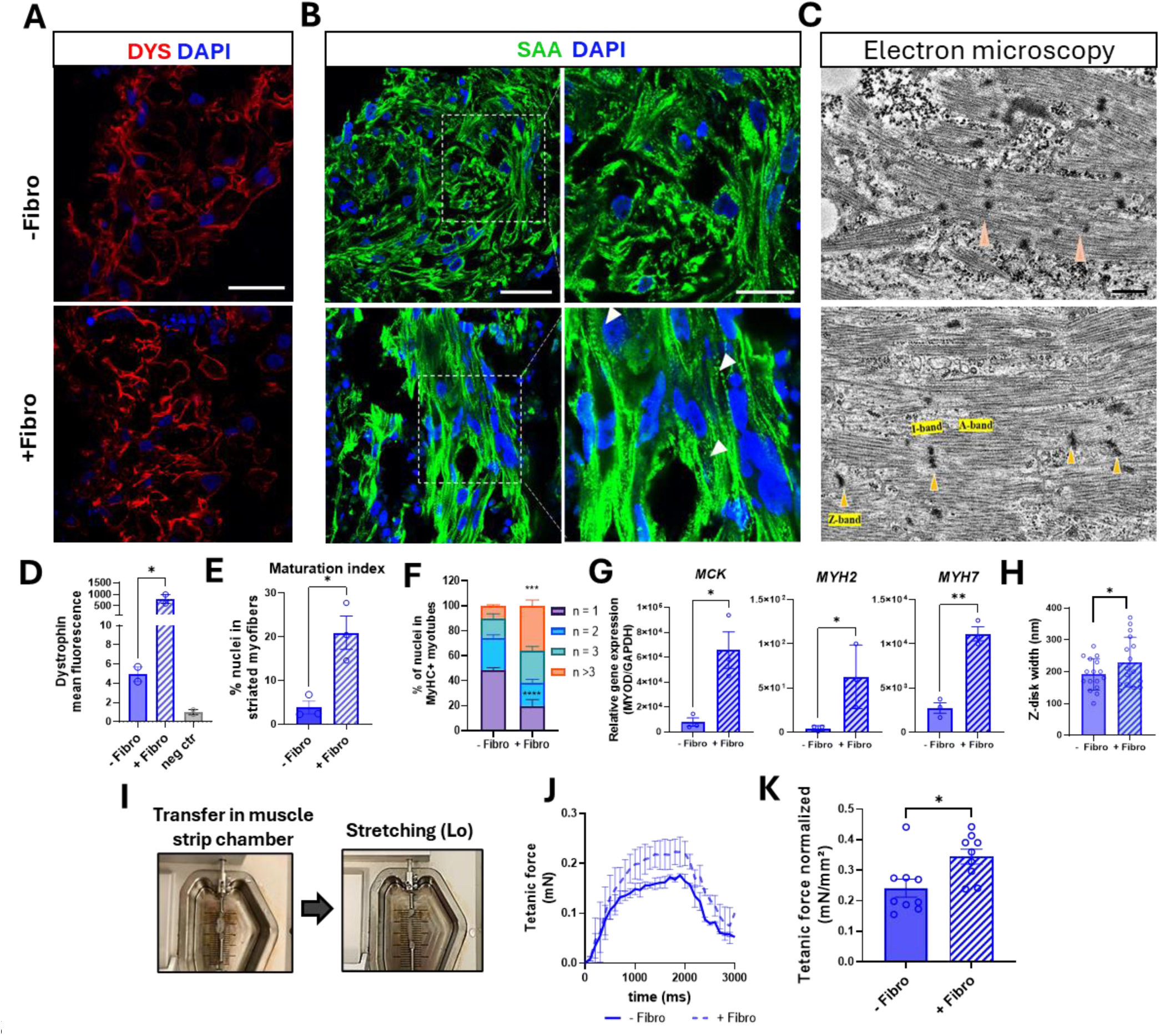
Structural and functional maturation in fibroblast-including MYOrganoids. **(A)** Representative transversal sections stained for Dystrophin (DYS). Scale bar: 40 µm. Nuclei were visualized with DAPI. (**B**) Representative longitudinal sections of MYOrganoids +/− fibroblasts (fibro), immunostained for Sarcomeric α-Actinin (SAA) and nuclei visualized with DAPI. Scale bars: 40 µm, enlargement 10 µm. (**C**) Transmission electron microscopy images showing sarcomeric structures. Orange arrows: Z-lines. Scale bar: 500 nm. (**D**) DYSTROPHIN staining quantification represented as mean intensity fluorescence and expressed as fold change to the negative control (sections stained without first antibody). N=2 (**E**) Quantification of maturation index was performed by calculating the nuclei inside striated myofibers visualized by SAA staining as a percentage of the total number of myofibers. N=3 (**F**) Quantification of fusion index calculated as % Myosin Heavy Chain positive (MYHC+) myotubes containing different numbers of nuclei (n) as depicted. N=3 (**G**) Gene expression analysis of MCK, MYH2 and MYH7, reported as gene expression relative to MYOD expressing population. N=3 (**H**) Width of Z-line visualized by Electron microscopy, in MYOrganoids with and without inclusion of fibroblasts. N=16 (**I**) Contractile muscle force analysis of MYOrganoids using a muscle-strip-based organ bath system. Lo, the optimal length used for normalization of force data (see methods). (**J**) Representative tetanic traces of MYOrganoids with and without the inclusion of fibroblasts. N=3 (**K**) Normalized tetanic force peak in MYOrganoids +/− fibroblasts. N=9). Data are presented as mean +/− SEM. Unpaired t-test was applied for statistical analysis (*p ≤0.05, **p ≤ 0.01).

We then assessed whether our MYOrganoids were functional by evaluating their physiological response to contraction stimulations, using a muscle organ bath system based on electrical pacing ^49^. To evaluate muscle force, MYOrganoids were transferred to the muscle strip chamber and stretched until the optimal length (Lo) for functional analysis (**Figure 2I**). Isometric force analysis revealed significantly higher tetanic force in MYOrganoids containing fibroblasts compared to the control (**Figure 2J**). Values were then normalized for the cross-sectional area (CSA) using the weight and optimal length of contraction established for each MYOrganoid ^50,51^ and expressed as specific tetanic force (mN/mm^2^) (**Figure 2K**). In particular, the highest force with fibroblasts had peak values ranging from 0.3 to 0.5 mN versus 0.1 to 0.2 mN in MYOrganoids without fibroblasts after normalization (**Figure 2K**). These data demonstrated that MYOrganoids plus fibroblasts have an improved structural organization and functional maturation that enables force contraction studies by electrical pacing.

### Inclusion of fibroblasts in DMD iPSC-derived MYOrganoids leads to increased muscle fatigue and pro-fibrotic signature

The improved muscle organization and functional maturation shown by MYOrganoids including fibroblasts, prompted us to exploit fibroblast features in disease modeling for DMD, where their role in disease progression is well known ^11,12^. We incorporated DMD fibroblasts to recapitulate the pathogenic microenvironment arising from their profibrotic activity exerted by tissue remodeling and matrix deposition ^12,52^. For that purpose, we used three DMD iPSC with different *DMD* mutations, a deletion of exon 45 (DMDdEx45), a deletion of exons 8-43 (DMDdEx8-43) and a deletion of exons 8-9 (DMDdEx8-9) with their isogenic control, the DMD dEx6-9 iPSC corrected to restore dystrophin expression ^53^ (hereafter called IsoCTR). We additionally used two control iPSC lines derived from healthy patients (CTR1, CTR2). Control and DMD MYOrganoids were generated from control and DMD iPSC with healthy or DMD human immortalized fibroblasts for functional and transcriptomic analysis (**Figure 3A**). The myogenic differentiation ability of both CTR and DMD iPSC lines was first evaluated in 2D cultures. All cell lines showed comparable myogenic potential, as expected using a direct myogenic conversion protocol bypassing any defective developmental steps (**Figure S2A**). Histological characterization in MYOrganoids showed the absence of dystrophin protein in DMD MYOrganoids and efficient myogenic differentiation and maturation in CTR and DMD MYOrganoids, as shown by the striated pattern visualized by SAA-stained sections, confirming the positive role of fibroblasts in enhancing maturation also in a DMD context (**Figure 3B**).

**Figure 3.**
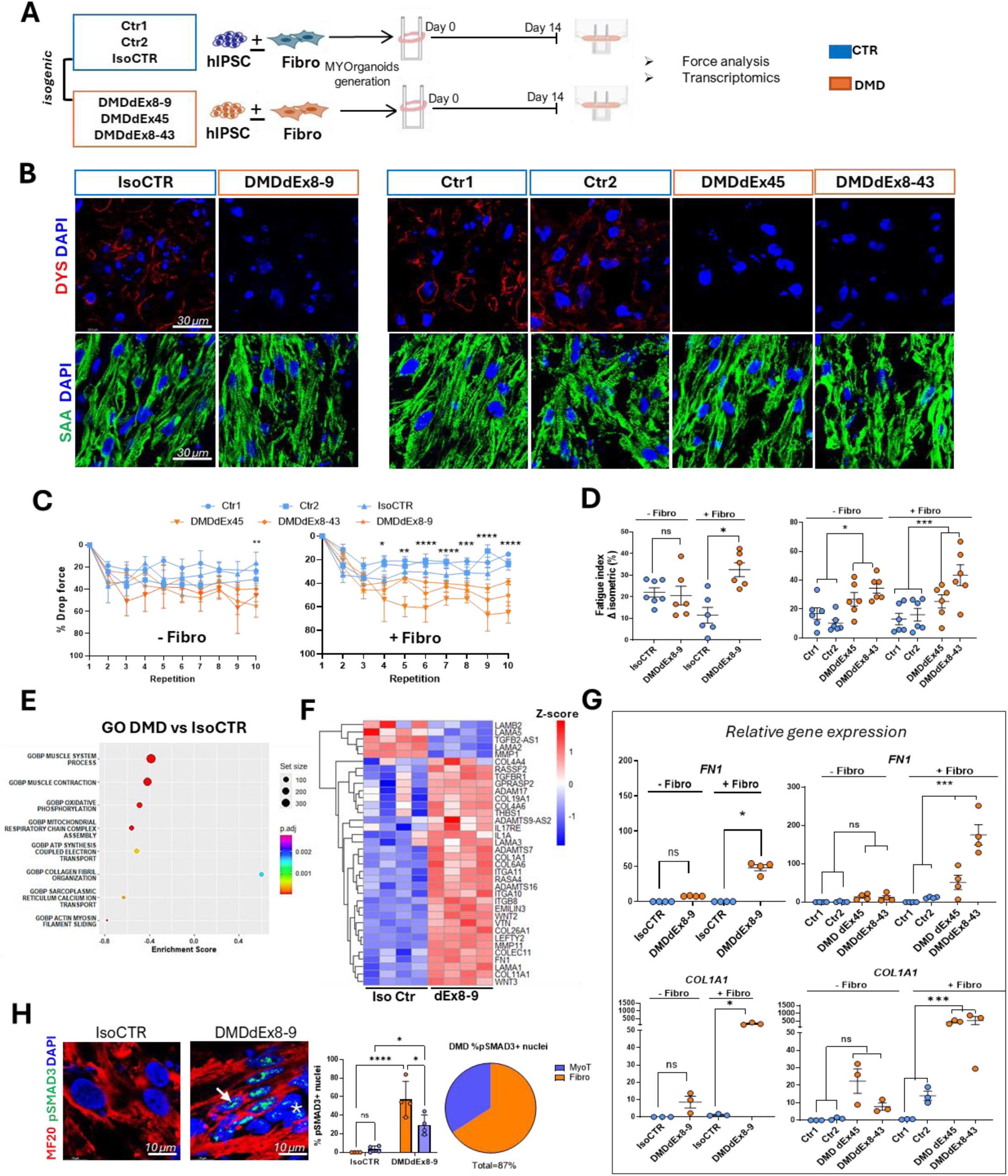
Fibroblast inclusion in DMD iPSC-MYOrganoids induces exacerbation of profibrotic signature and fatiguability. **(A)** Overview of MYOrganoids generation from control iPSC (CTR1 and CTR2), DMD iPSC (DMDdEx45, DMDdEx8-43) and isogenic iPSC (IsoCTR and DMDdEx8-9) including control or dystrophic fibroblasts respectively for histological characterization, force and transcriptomic analysis**. (B)** Immunostaining of MYOrganoids cross sections for Dystrophin (Dys), Sarcomeric a-actinin (SAA), and DAPI for nuclei. Scale bar: 30 µm **(C)** Drop force over 10 repetitions of eccentric (ECC) contractions in all CTR and all DMD MYOrganoids without (−) or with (+) fibroblasts N=5. **(D)** Fatigue index calculated as % of drop force between two isometric contractions (ISO) performed before and after 10 repetitions of ECC in all CTR and all DMD MYOrganoids +/− respective fibroblasts. N=6. **E)** Gene Ontology enrichment analysis (GO TERM) from RNA-seq data in isogenic iPSC-derived CTR (IsoCTR) and DMD (DMDdEx8-9). The scale of color is representative of the p adjusted (p.adj) N=4 **F**) Clustered heatmap of fibrotic Differentially Expressed Genes (DEG) relative to DMD vs IsoCTR (left). Color scale represents the Z-score (red is higher, blue lower) **G)** Relative gene expression of Fibronectin (FN1) and Collagen (COL1A1) in all iPSC CTR and DMD-derived MYOrganoids with or without fibroblasts. For FN1, N=4 and for COL1A1 N=3 **H)** Representative images showing activation of TGF-β signaling by immunostaining for pSMAD3+ cells in MyHC+ myotubes (arrows), and in fibroblasts (asterisk), considered as MyHC-cells, and relative quantification. Scale bar: 10 µm, N=4 For all panel, data are presented as mean ± SEM. For panel A-G, unpaired one-tailed t-test was performed for statistical purposes. For panel H, a 2-way ANOVA test was performed (F=1,12, degree of freedom=12) (*p ≤0.05, **p ≤ 0.01, ***p ≤ 0.001, ****p ≤ 0.0001, ns= not significant).

To assess whether DMD MYOrganoids display hallmarks of DMD pathophysiology, we evaluated muscle function, which represents one of the most difficult challenges in establishing therapeutic readouts with *in vitro* systems. To identify reliable force parameters reflecting the defective DMD muscle performance, MYOrganoids were subjected to isometric contractions to measure tetanic force, and to eccentric contractions, which play a critical role in the disease progression of DMD^54^ by triggering the membrane degeneration/regeneration cycles^55^ to assess muscle strength and fatiguability. No significant changes were observed in the tetanic isometric force between CTR and DMD MYOrganoids (**Figure S2B**), as expected for muscles never been challenged by contractions and with the same myogenic potential, while DMD MYOrganoids showed higher drop force in the eccentric contraction repetitions exclusively when including DMD fibroblasts starting from the 4^th^ eccentric repetition (**Figure 3C**). On the other side, CTR and DMD MYOrganoids not including fibroblasts, show significant differences in drop force just at the tenth repetition (**Figure 3C**). To accurately quantify muscle fatigue, we calculated the fatigue index as the drop of force between the isometric contractions performed before and after the 10 repetitions of eccentric exercise. The analysis showed a significantly higher fatigue index in DMD MYOrganoids as compared to CTR MYOrganoids and remarkably, this phenomenon was highly accentuated by the presence of fibroblasts. (**Figure 3D**). To check whether the impact on disease exacerbation and increase of difference in fatiguability between CTR and DMD MYOtissue was affected by the different genetic backgrounds of the fibroblast source, we employed three additional fibroblast cell lines from either healthy and DMD individuals to generate CTR and DMD MYOrganoids. Muscle force analysis confirmed that the fatigue index in DMD MYOrganoids was significantly higher than CTR MYOrganoids, regardless of the genetic source of either control or DMD fibroblasts. This analysis also confirmed the display of higher and significant muscle fatiguability in fibroblasts-including MYOrganoids (**Figure S2C**). Collectively, these data indicate that DMD MYOrganoids including DMD fibroblasts, through their pro-fibrotic activity, display exacerbated loss of muscle resistance and increase in fatiguability (**Figure 3A-D**). The finding also demonstrates that eccentric-based drop force evaluation is a meaningful and reliable therapeutic readout, as significant differences were detected between CTR and DMD MYOrganoids including fibroblasts.

We then performed transcriptomic analysis in the isogenic iPSC lines (IsoCTR and DMDdEx8-9) to better characterize the pathogenic hallmarks introduced by fibroblast incorporation within the organoids. Analysis of significant (p.adj < 0.05) and relevantly different transcripts (abs log2FoldChange > 1) revealed 4610 differentially expressed genes (DEGs) between DMDdEx8-9 and IsoCTR MYOrganoids, of which 2181 upregulated and 2429 downregulated (**Figure S3A-B**). Of those, we identified genes involved in decreased muscular contraction (e.g. muscle system process, muscle contraction, sarcoplasmic reticulum calcium ion transport and actin-myosin filament sliding) as well as reduced energetic molecular process (e.g. oxidative phosphorylation, mitochondrial respiratory chain complex assembly, ATP synthesis coupled electron transport), as depicted by gene ontology biological process (GOBP) enrichment analysis (**Figure 3E, table S1**). Remarkably, between the upregulated pathways in DMD MYOrganoids, we identified higher collagen fibril organization with an enrichment score of +0.6 (**Figure 3E**). This GO analysis was also confirmed by Kyoto encyclopedia of genes and genome (KEGG) enrichment analysis, which showed a downregulation of oxidative phosphorylation and an upregulation of ECM receptor interaction, such as Integrin alpha and beta (**Figure S3C**). Interestingly, the analysis of genes involved in the ECM remodeling revealed an upregulation of genes coding for collagens, laminins, metalloproteases and TGF-β related genes such as TGF-B receptor (TGFBR1) and the ncRNA inhibitor of TGF-β (TGFB2-AS1) in DMD MYOrganoids compared to isogenic control (**Figure 3F**). Additionally, we found increased levels of latent TGF-β binding proteins (LTBP 2 and 4) in dystrophic MYOrganoids, indicating that TGF-β is activated in the DMD context and contributes to ECM remodeling (**Table S1**).

The increased profibrotic signature in DMD iPSC-derived MYOrganoids was then confirmed in all CTR and DMD iPSC-derived MYOrganoids, by evaluation of the expression of two key fibrotic markers, Fibronectin-1 (*FN1*) and Collagen-1 (*COL1A1*). Importantly, the differences in the expression of fibrotic markers between CTR and DMD MYOrganoids were significant only when including fibroblasts (**Figure 3G**). We further examined, at histological level, the presence of activated TGF-β signaling in DMD organoids by looking at phosphorylated SMAD3 (pSMAD3), the transcriptional effector of the canonical TGF-β pathway. Notably, DMDdEx8-9-derived MYOrganoids showed increased pSMAD3 positive nuclei both in myotubes (MyHC positive staining) and fibroblasts (MyHC negative cells) (**Figure 3H**). This observation indicates a crosstalk between fibroblasts and myofibers and supports the potential of using fibroblasts in tissue engineering as a source of ECM and pro-fibrotic cues under pathological conditions.

### Juxtracrine role of fibroblasts in the modeling of DMD phenotype

We then sought to gain insights into the contribution of fibroblasts in the recapitulation of key pathogenic hallmarks of DMD. Given that fibroblasts are known to primarily act through their secreted molecules ^56–58^ we analyzed the secretome of both control (CTR) and DMD MYOrganoids (MyoT + Fibro) and compared it to that of single-cell 3D cultures (Fibro-only and MyoT-only) (**Figure 4A**). Analysis of culture media on day 14 showed elevated levels of secreted TGF-β and Collagen 4 (COL4) in DMD MYOrganoids cocultured with fibroblasts compared to control conditions. No significant differences were observed between control and DMD in Fibro-only and MyoT-only 3D cultures (**Figure 4B**). These data indicate that DMD MYOrganoids in coculture with fibroblast exhibit an increased release of fibrotic factors.

**Figure 4.**
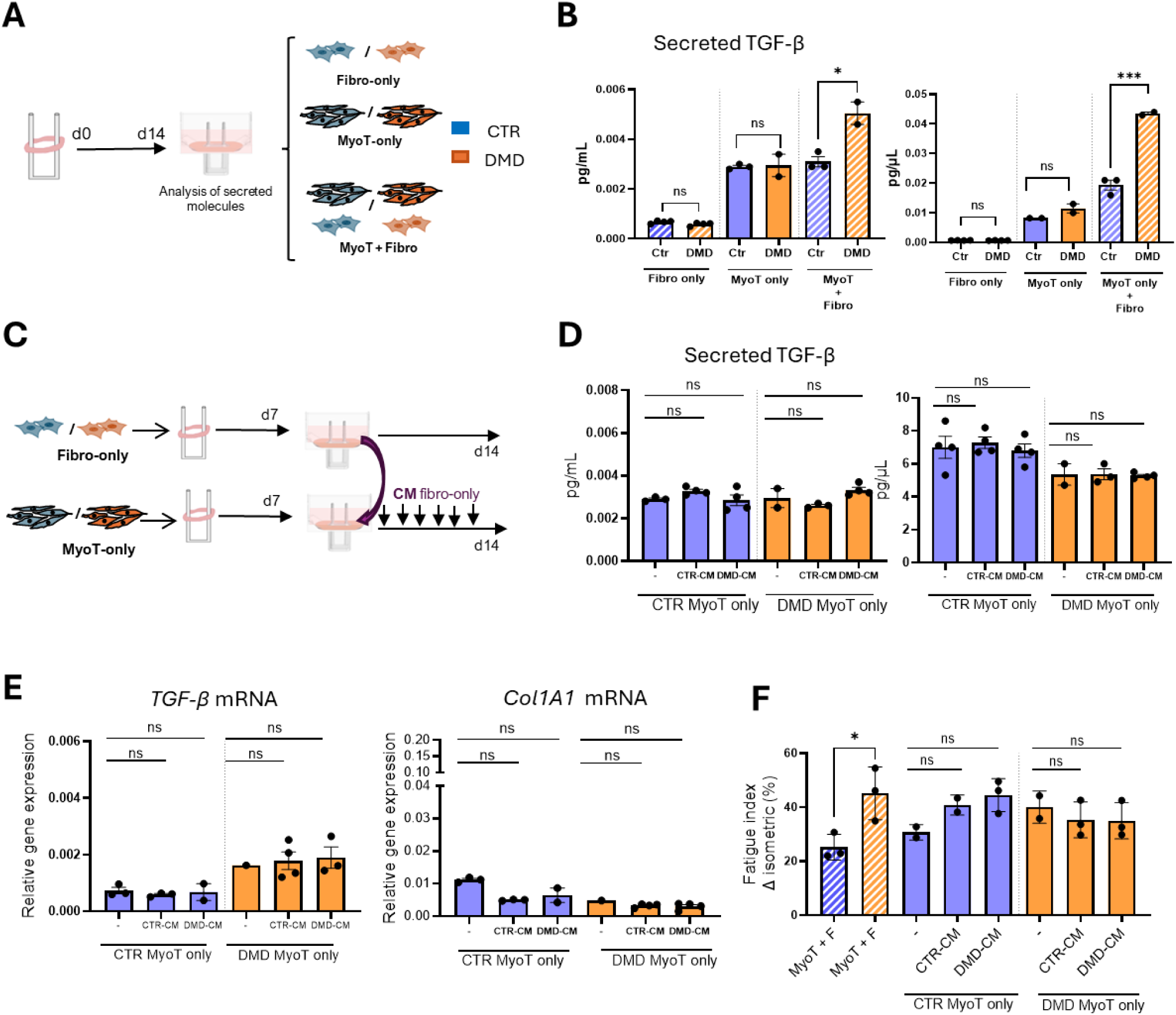
Fibroblast role in the modeling of DMD phenotype requires direct contact with DMD iPSC-derived muscle cells. **A)** Scheme of engineered 3D cultures created using either iPS-derived myotubes (MyoT), fibroblasts (Fibro), or a combination of both (MyoT + Fibro=MYOrganoids) from CTR or DMD cells. **B)** TGF-β and COL1A1 ELISA performed in Fibro only, MyoT only or combination (Fibro+MyoT). N=3 **C)** Scheme of conditioned medium (CM) exchange. From day 7, (CM) was collected daily from 3D cultures with fibroblasts (Fibro) alone and used daily in MyoT 3D cultures compared to untreated MyoT only or MyoT+Fibro co-cultured MYOrganoids.N=3 **D)** TGF-β and Fibronectin 1 (FN1) levels detected by ELISA in CTR or DMD only MyoT only organoids after 7 days of treatment with CM from either wild-type fibroblasts (CTR-CM) or DMD fibroblasts (DMD-CM) and analyzed at day 14.N=3 **E)** TGF-β and COL1A1 gene expression in in CTR or DMD only MyoT only organoids after 7 days of treatment with CM from either wild-type fibroblasts (CTR-CM) or DMD fibroblasts (DMD-CM) and analyzed at day 14. N=3 **F)** Fatigue index in MyoT + Fibro organoids, and in MyoT only tissues treated or not with CTR-CM and DMD-CM. CTR conditions are indicated in blue and DMD organoids in orange. N=3. For all panels, data are represented as average ± SEM. *p < 0.05; ***p<0.0001; ns = not significant (unpaired one-tailed Student’s t test).

To determine whether fibroblasts amplify the fibrotic phenotype through their secretory activity, we collected conditioned media (CM) from control (CTR) and DMD Fibro-only 3D cultures on day 7 of *in vitro* growth. We then applied this CM to treat CTR and DMD MyoT-only tissues for additional 7 days, assessing the secretion of fibrotic factors, gene expression, and muscle force (**Figure 4C**). ELISA assay on secreted TGF-β and fibronectin 1 (FN1) revealed no difference in MyoT-only 3D cultures treated with the CM from either CTR or DMD Fibro-only tissues (**Figure 4D**). Congruently, gene expression levels of TGF-β and Col1A1 were similar in CTR and DMD MyoT-only cultures, regardless of the treatment with the CM (**Figure 4E**), while those were higher just in DMD cocultures (MyoT + Fibro) compared to control conditions (**Figure S4A**). The lack of effect of the CM indicates that the profibrotic increase seen in DMD MYOrganoids in coculture with fibroblasts occurs by a juxtracrine effect, as it comes from contacts between the two cell types.

We further evaluated the requirement of fibroblasts-myotubes coculture in disease exacerbation by assessing muscle force analysis following CM treatment, which confirmed no difference in fatiguability in myotube-only 3D cultures treated with control or DMD Fibro-derived CM, unlike higher fatigue index in DMD organoids containing both fibroblasts and myotubes in coculture (**Figure 4F**). Interestingly, MYOrganoids including fibroblasts displayed a higher tetanic force compared to myotubes-only cultures treated or not with fibroblasts CM, also supporting their cell contact-dependent role for enhanced functional maturation (**Figure S4B**). Collectively, analysis of secreted protein and force under CM experiments showed that the dystrophic fatiguability and fibrosis traits emerged only when fibroblasts and muscle cells were cocultured, indicating that contact-dependent cellular communication between muscle and fibroblasts is required to reveal the DMD traits of profibrotic secretion and fatiguability.

### AAV-microdystrophin gene transfer rescues muscle resistance while partially restoring Dystrophin Glycoprotein Complex components in DMD MYOrganoids

As proof of concept that MYOrganoids were suitable as a screening platform for gene therapy, we used AAV-mediated delivery of microdystrophin (µDys) and assessed its therapeutic efficacy in the DMD context. We used AAV9 capsid for this aim, as this viral serotype was used in recent clinical trials and showed a high transduction rate in patients’ myofibers ^59,60^. We infected the organoids with a codon-optimized µDys gene (dR4-23, same protein product used in clinical trials ^61^) under the control of the muscle-specific spc512 promoter ^16^ for gene transfer using AAV9 (AAV9-µDys) in all DMD MYOrganoids iPSC (dEx45, dEx8-43 and dEx8-9 with isogenic control, IsoCTR) and assessed gene transfer efficiency, muscle force analysis and membrane stability assessment (**Figure 5A**). We first optimized the infection conditions using the reporter AAV9-CMV-GFP in CTR MYOrganoids including fibroblasts. Infection was performed on day 7 of the differentiation protocol, diluting AAV particles directly in the medium, and maintained for an additional 7 days (**Figures S5A-C**). Optimal low and high AAV9-µDys doses were established based on previous dosing studies to have intermediate transduction levels at low doses (1E+09 vg/MYOrganoid), and high transduction levels at high doses (5E+10 vg/MYOrganoid) (**Figure S5D**). Gene transfer efficiency was evaluated by quantification of viral copy number (VCN) on genomic DNA and by the expression level of the transgene and the encoded protein, showing a clear dose-dependent entry and expression of µDys in DMD MYOrganoids (**Figure 5B, S6A-B**). As expected by using a muscle-specific promoter, only the muscle cells (MyoT) but not the fibroblasts expressed the transgene after infection of AAV9-uDys in the single populations (**Figure S6C**). We then tested whether µDys affected the contractility and fatigue resistance of the infected organoids. Isometric tetanic force analysis did not reveal any significant changes in DMD MYOrganoids following µDys delivery compared to not-infected conditions (**Figure S6D**), as expected by first-time contracting muscle tissues. We then challenged the muscles with repeated eccentric repetitions, and we observed a dose-dependent attenuation of the drop force in DMD MYOrganoids after µDys infection (**Figure 5C**). Consistently, the fatigue index was greatly reduced in a dose-dependent manner in all DMD MYOrganoids treated with a high dose of µDys, although reaching significance only for DMDdEx45 and DMDdEx8-9 MYOrganoids (**Figure 5D**). To confirm that the observed effects were specifically due to µDys expression rather than AAV infection alone, we infected DMD MYOrganoids with a control vector containing a dystrophin fragment lacking a promoter (hereafter called empty vector), which infection did not result in protein expression (**Figure S6E-G**). Particularly, the MYOrganoids infected with the empty vector did not show any differences in the fatigue index analysis, at both low and high dose, while we observed a decreased fatigue index in DMD MYOrganoids infected with an AAV containing µDys (**Figure S6H**). Overall, the muscle force data confirmed that ectopic µDys expression rescues muscle endurance and fatiguability.

**Figure 5.**
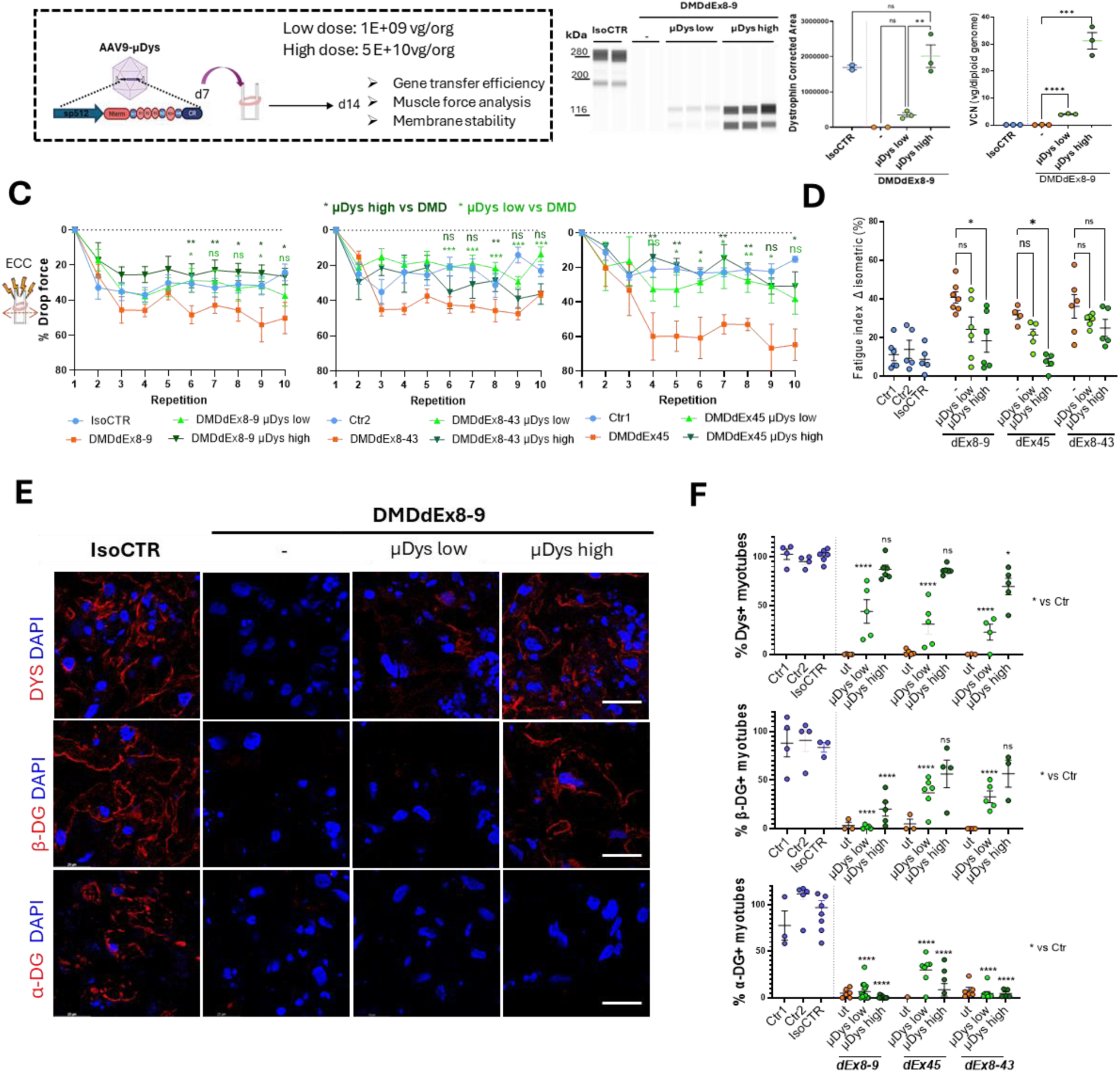
µDys gene transfer-mediated improvement of muscle resistance and partial rescue of DCG complex. (**A**) Scheme of AAV9-µDys infection and doses used in DMD iPSC-derived MYOrganoids. (**B**) Evaluation of gene transfer efficiency by viral copy number (VCN) analysis (N=3) and protein expression analysis by capillary western blot in isogenic iPSC DMD dEx8-9-derived MYOrganoids infected with low and high dose of µDys versus non-infected (−) IsoCTR (N=3). **(C)** Drop force over 10 repetitions of eccentric (ECC) contractions in isogenic cell lines and CTR1, CTR2 and DMDdEx45, DMDdEx8-43-derived MYOrganoids receiving low and high doses of µDys. N=6 **(D)** Fatigue index in all CTR and DMD MYOrganoids treated or not with m-Dystrophin at low and high dose. N=5. **(E)** Histological evaluation of Dystrophin Glycoprotein Complex (DGC) upon infection, by immunostaining for Dystrophin (DYS), α- Dystroglycan (α-DG) and β- Dystroglycan (β**-**DG) in cross sections of Isogenic CTR and DMD-iPSC-derived MYOrganoids. Scale bar: 40 µm and (**F**) relative quantification in all control and iPSC-derived MYOrganoids N=4.. Data are presented as means +/− SEM. Statistical analysis was performed with an ordinary one-way ANOVA test (*p ≤ 0.05, **p ≤ 0.01, ***p ≤ 0.001, ****p ≤ 0.0001, ns = not significant) with multiple comparisons corrected with Dunnett’s test (F=3,24 and degree of freedom = 24)

Because dystrophin exerts its biomechanical support by holding the Dystrophin Glycoprotein Complex (DGC) at the membrane, we monitored key components of this complex, such as the transmembrane β- Dystroglycan (β-DG), directly binding Dystrophin, and the extra-cellular α-Dystroglycan (α-DG), whose proper expression and localization are impaired in the absence of dystrophin ^62^. To this aim, DMD MYOrganoids infected with low and high dose of µDys, were subjected to histological analysis. Quantification of µDys showed around 25-30% of dystrophin-positive myotubes at low dose compared to 85% in the high dose condition in DMD dEx8-9 (**Figure 5E-F**) and dEx45 and 65% in the DMD dEx8-43 MYOrganoids (**Figure S7A-B**). Immunostaining on transversal MYOrganoids showed a dose-dependent yet not complete restoration of β-DG in all DMD iPSC-derived MYOrganoids. Interestingly, even high doses ensuring nearly total Dystrophin transduction in all the DMD MYOrganoids did not significantly restore α-DG (around 10% in DMDdEx8-9 and DMDdEx8-43 and around 30% in DMDdEx45) suggesting a potential therapeutic limitation of µDys (**Figure 5E-F**). Importantly, these data demonstrate the limited restoration of Dystrophin-associated components to the sarcolemma after successful µDys gene transfer.

### Optimal microdystrophin gene transfer partially corrects the transcriptomic profile of DMD MYOrganoids and does not rescue the profibrotic signature

To examine the effect of microdystrophin (µDys) on the signaling pathways and molecular processes characteristics of dystrophic pathology, we performed bulk RNA-seq analysis comparing the corrected DMD dEx8-9 (IsoCTR) and DMDdEx8-9 MYOrganoids untreated and treated with a high dose of AAV9-µDys (n=4).

Principal component analysis (PCA) of the top 10000 genes with the highest variance showed a distinct transcriptomic profile of DMD organoids treated with µDys (dEx8-9 + µDys high) and untreated (dEx8-9) compared to isogenic controls (IsoCTR) (**Figure S8A**). To evaluate the effect of µDys in restoring key pathways involved in DMD, we firstly compared the transcriptomic profile of µDys-treated DMD organoids to untreated DMD organoids (dEx8-9 µDys vs dEx8-9), and then to the isogenic control (dEx8-9 µDys vs IsoCTR). Particularly, µDys-treated DMD organoids showed a total of 327 DEGs, of which 154 upregulated and 173 downregulated, compared to untreated organoids (**Figure 6A, Table S2**). Of those 327 DEGs, 52% (170/327) are in common with the DEGs that were found dysregulated in DMD organoids when compared to the isogenic control (**Figure S8B**). Therefore, we performed the gene ontology enrichment analysis of the common category to see the effect of μDys on the DMD-dysregulated pathways. This analysis revealed increased expression of genes involved in the microtubule organization (GOBP: Microtubule organizing center organization NES = +2.31), cytoskeleton organization (GOBP: Protein localization to cytoskeleton NES=+1.99), and a decreased expression of genes involved in the inflammation (GOBP: Regulation of inflammatory response, NES = −1.45) (**Figure 6B**).

**Figure 6:**
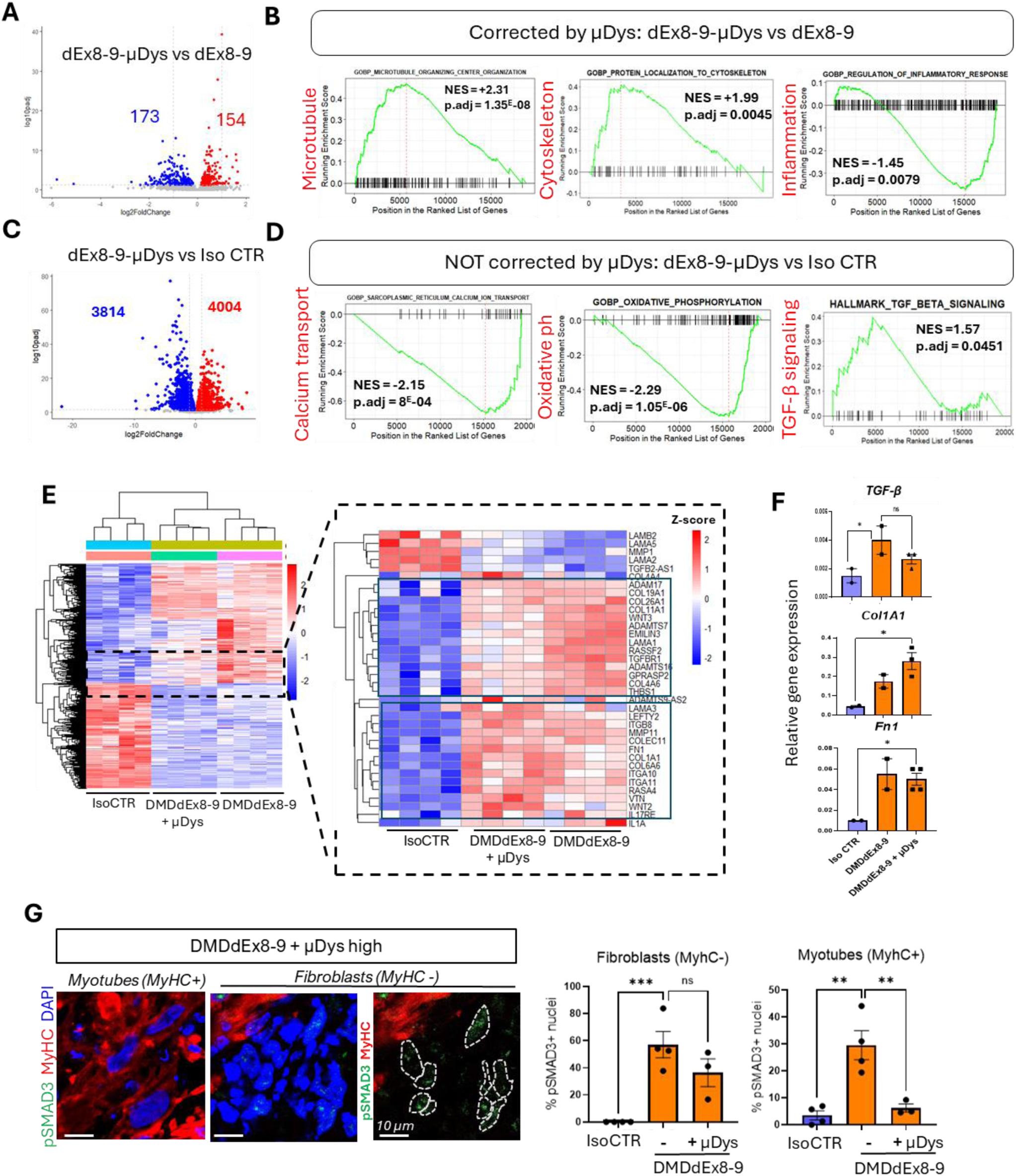
Persistent fibrotic activity in µDys-treated isogenic DMD MYOrganoids. **(A,C)** Volcano plot showing the differential gene expression in treated (dEx8-9-µDys) versus untreated (dEx8-9) DMD MYOrganoids(A) or DMD dEx8-9-µDys vs isogenic control (IsoCTR (C). Upregulated genes are shown in red, while downregulated genes are shown in blue. Numbers indicate the amount of downregulated and upregulated genes. **(B,D)** Gene ontology enrichment analysis of treated (dEx8-9-µDys) versus untreated (dEx8-9) DMD MYOrganoids (B) or dEx8-9-µDys vs IsoCTR (D). Normalized enrichment score (NES) and adjusted p-value (p.adj) are indicated for each subcategory. (**E)** Clustered Heat map and enlargement depicting relative transcript levels of differentially expressed genes in corrected DMD (IsoCTR) and DMD MYOrganoids transcriptomics treated or not with µDystrophin (dEx8-9 and dEx8-9 µDys high). Lower and higher expressions are depicted in blue and red, respectively. The data points above the significance threshold (p adjusted < 0.05, fold2change > 2) are marked in blue (downregulated) and red (upregulated), and others are marked in gray (not significant). (**F**) Relative gene expression of TGF-β, ColA1 and FN1 in Isogenic control, DMD dEx8-9 untreated or tretated with μDys. (**G**) Histological staining of phospho-SMAD3 (pSMAD3) staining in green, myosin heavy chain (MyHC) in red and nuclei with DAPI in µDys-treated DMD MYOrganoids, and relative quantification of pSMAD3 positive nuclei in percentage. Scale bar = 10 µm. For panel F-G, data are presented as means +/− SEM. For panel F, statistical analysis was performed with an ordinary one-way ANOVA test (*p ≤0.05, ns = not significant), while for panel G, unpaired one-tailed Student’s t test was performed (** p < 0,005; *** p < 0,001; ns = not significant).

We then analyzed the DEGs between the treated organoids and the isogenic control to check the transcriptomic signature not restored by µDys (dEx8-9-µDys vs IsoCTR). Particularly, µDys-treated organoids present 7818 DEGs (4004 upregulated and 3814 downregulated) when compared to the corrected isogenic organoids (**Figure 6C**). Gene ontology analysis of the DEGs between µDys-treated DMD organoids and IsoCTR organoids revealed the pathways that are not restored by the gene transfer. Particularly, treated organoids showed persistent downregulation of genes involved in calcium transport (GOBP: Sarcoplasmic reticulum calcium ion transport, NES = −2.15), oxidative phosphorylation (GOBP: Oxidative phosphorylation, NES = −2.29) and upregulation of TGFβ signaling (Hallmark TGFβ signaling, NES = +1.57), when compared to isogenic controls (**Figure 6D, Table S3**). Consistently, a clustered heatmap of all 1716 most significant differentially expressed genes (DEGs) further highlighted the transcriptomic differences between control and DMD organoids (**Figure 6E**).

Analysis of Z-score of fibrotic genes among the three conditions (IsoCTR, DMDdEx8-9 and DMDdEx8-9 treated with µDys) revealed a persistent but variegated profibrotic signature. In particular, we revealed a slight decrease in the expression of ADAMs (i.e. ADAM17, ADAMTS7, ADAMTS16), collagen genes (e.g. COL11A1, COL4A6) and the TGF-β receptor 1 gene (TGFBR1) in µDys-treated DMD organoids, when compared to the untreated condition, although still significantly different from isogenic control (**Figure 6E, right panel).** Additionally, other fibrotic genes, like *FN1* and *COL1A*1, Integrins or cytokines (e.g.IL1A), showed no expression changes compared to untreated control (**Figure 6E, right panel**). Further validation of the expression of *COL1A1*, *FN1,* not decreased by the µDys treatment, and of *TGF-β*, partially downregulated, confirmed the persistent expression of those genes in DMD-treated MYOrganoids (**Figure 6F**). To understand the mechanisms behind the unresolved fibrotic process, we evaluated the reduction in TGF-β pathway activation within both cell subpopulations in DMD MYOrganoids following µDys treatment. Histological staining revealed a significant decrease in pSMAD3-positive nuclei within the skeletal muscle subpopulation of µDys-treated organoids, whereas no difference was observed in fibroblasts after µDys treatment (**Figure 6G**). This result suggests a cell-autonomous correction of this signaling within the muscle population receiving µDys, but no paracrine effect on fibroblast activity. The persistence of activated TGF-β signaling in the fibroblast subpopulation after µDys gene transfer (**Figures 3H and 6G**) can therefore explain the uncorrected profibrotic signature and poses the basis for future investigation aimed at improving or driving beneficial cell-cell communication.

## Discussion

Here, we present a novel human-relevant model that simulates key traits of advanced stages, enabling the interrogation of gene replacement effect on muscle function and fibrosis, and facilitating the identification of new therapeutic avenues.

We report the generation of 3D-engineered muscles called MYOrganoids, composed of a homogeneous population of iPSC-derived skeletal muscle cells ^35,40^ and fibroblasts, major players in muscle tissue organization and microenvironment regulation^37,38,63^. We proved that fibroblast inclusion in the MYOrganoids results in improved structural maturation that allows high functional performance under muscle force evaluation, while also eliciting a fibrotic signature in a DMD context.

Importantly, fibroblast incorporation was pivotal in exacerbating DMD phenotype and revealing fatiguability, which is a top hallmark for DMD muscle function, thereby enabling its unprecedented use as therapeutic readouts in vitro. This is important considering that the capacity to evaluate muscle function in a high throughput manner remains limited, further hindering progress in testing or developing effective treatments. As such, while several remarkable studies reported on muscle force in 3D models ^32,33,53,64,65^, evaluating muscle function still presents challenges due to the complexity of identifying disease-specific force parameters and the variability of *in vitro* models.

Furthermore, our study sheds new light on the mechanism underlying the contribution of fibroblasts to the modeling of DMD phenotype. We showed that direct interaction of DMD fibroblast with muscle cells is essential to reproducing key pathogenic hallmarks, such as a profibrotic signature and increased muscle fatigue in DMD MYOrganoids. Interestingly, fibroblast role in the recapitulation of DMD hallmarks was not exerted by their profibrotic secretory activity per se but required direct interaction with iPSC-derived muscle cells. As such, fibroblasts in a DMD context showed higher secretion of fibrotic factors like TGF-β, fibronectin, and collagen through cell-contact signaling, suggesting that cell communications between the two cell populations, fibroblasts and muscle fibers, is crucial to reveal pathogenic traits. One possible hypothesis is that fibroblasts secretory activity is enhanced by alterations in cell communications with dystrophin-deficient muscle cells whose membrane stability is impaired, thereby causing increased matrix deposition and susceptibility to contraction-induced damage and fatigue.

Overall, the exacerbated severity of DMD MYOrganoids enabled the evaluation of the therapeutic potential of µDys, currently employed in clinical trials. Consistently with the partial efficacy in the clinics, µDys ^16,17^ delivery in DMD MYOrganoids did not fully rescue DMD phenotype as it led to incomplete restoration of the components of the dystroglycan complex and no reduction in fibrotic genes transcriptome. Notably, we observed a reduction in the TGF-β pathway activation exclusively on muscle cells, while no effect was observed in fibroblasts. This can explain the persistence of ECM protein production and secretion in µDys treated DMD MYOrganoids, while we observed an effect in the restoration of muscle strength.

Our findings align with previous results showing that μDys exhibits limited effectiveness in reducing fibrosis in advanced disease stages ^66^ and in more severe models and muscles affected by the disease, such as the diaphragm in the Dba2 *mdx* model ^67^. We can speculate that the partial restoration of DAG complex after gene transfer causes mechanical alterations of the membrane that are translated as profibrotic signals. In advanced disease stages, the stiffened ECM perpetuates fibrosis through mechanotransduction, activating fibroblasts to deposit ECM proteins and increasing rigidity. This feedback loop hinders microdystrophin’s ability to reverse fibrosis.

Up to now, no other work revealed the impact of µDys on the fibroblast population of the muscle, revealing the potential of our system in highlighting specific cell-cell communications in response to treatment. Further work is needed to fully understand the role of fibroblasts in the dystrophic process. For example, how the absence of dystrophin in the myotubes is influencing fibroblast activity, and how the reintroduction of µDys is affecting the composition of the ECM, still needs to be elucidated.

Overall, pro-fibrotic DMD MYOrganoids provide a valuable tool either for the investigation of mechanisms driving dystrophic process and as a human *in vitro* counterpart to animal *in vivo* preclinical studies, with the potential to unravel therapeutic limitations of current DMD treatments and accelerate the identifications of new treatments.

## Acknowledgments

We express our gratitude to Rene Hummel (Danish MyoTechnology), Guillaume Tanniou, and Nicolas Guerchet for their technical support in muscle force evaluation. We thank Guillaume Corre for the aid with RNA-seq analysis. We thank the service of bioproduction of Genethon, for preparing the AAV particles used in this study. We also extend our appreciation to Jérémie Cosette for the assistance in imaging and to the histology team for their technical expertise. Finally, we would like to thank Frederique Magdinier for generously providing the DMD iPSC dEx45 and CTR1 iPSC.

## Author contributions

SA and LPa designed the experiments. LPa performed most of the experiments, including iPSC culture, organoid generation, imaging, and muscle force analysis. LPi performed force assays and gene and protein expression analysis. AJ performed secretome analysis. MM contributed to muscle force analysis. AVH analyzed RNA-seq data. RE and GB performed electron microscopy analysis. AB generated and provided the immortalized human fibroblasts. SA and LPa performed data analysis and figure preparation. SA conceived, supervised the project, and wrote the manuscript. All authors discussed the results. SA, LPa, DI and IR reviewed and edited the manuscript.

## RESOURCE AVAILABILITY

### Lead contact

Further information and requests for resources and reagents should be directed to and will be fulfilled by the lead contact, Sonia Albini.

### Material availability

CTR2, IsoCTR, DMDdEx8-9 and DMDdEx8-43 hIPS cell lines used in this study are restricted due to MTAs (Ref: MTA 2022 GENETHON - AIM _DMD, MMTA202109-0107, 2021-0296_MTA_CECS). This study did not generate new unique reagents.

### Data and code availability

The data supporting the findings of this study are available within the article and its supplemental information. The RNA sequencing datasets generated in this study can be found in the NCBI Bioproject database (https://www.ncbi.nlm.nih.gov/bioproject) using the access number PRJNA1208956.

## METHODS

### Cell culture

Human iPSCs used in the study were as follows: CTR1 and CTR2 from healthy individuals, DMDdEx45, DMDdEx8-43 from DMD patients. The isogenic iPSC DMDdEx8-9 and corrected control (CRISPR-mediated skipping of mutated exons dEx6-9, isogenic control) were obtained from Olson lab. The CTR1 and DMDdEx45 hiPSC lines were generated from skin fibroblasts from Coriell and were obtained from Marseille Stem Cells platform in Marseille Medical Genetics laboratory ^68^. iPSCs were maintained using mTeSRplus medium (Stem Cell Technologies) and passaged using ReleSR (STEMCELL Technologies) on Matrigel-coated wells (Corning). iPSC engineering and muscle differentiation were performed by adapting a transgene-based method previously described ^35,36^. All iPSC lines are engineered for transgene-based myotubes differentiation. Briefly, cells were nucleofected (4D Nucleofector, Lonza) with two enhanced versions of piggyBAC (ePB) containing respectively MyoD and BAF60c genes under tetracycline responsive promoter ^35,36^. Additionally, iPSCs were regularly selected for the presence of the two ePBs with puromycin (10 µg/ml) and blasticidin (20 µg/ml) at the same time (Sigma-Aldrich). Human immortalized fibroblasts from control and DMD patient were generated and obtained from Myobank-AFM of Myology Institute from muscle biopsies. Control and DMD human immortalized fibroblasts, used in co-culture with hiPSC in 3D MYOrganoids, were maintained in culture in DMEM High Glucose Lglut supplemented with 20% fetal bovine serum (FBS) and passaged using TrypleE (Gibco). Human primary fibroblasts from control DMD patients were obtained from DNA bank of Genethon Fibroblasts were maintained in DMEM high glucose (Gibco) supplemented with 20 ng/ml recombinant human fibroblast growth factor (h-FGF) (Peprotech) and 20% fetal bovine serum (Gibco). Primary fibroblasts were passaged using TrypleE (Gibco). All cell lines are listed in **table S4.**

### Generation of MYOrganoids

MYOrganoids from iPSCs were generated adapting the protocol described for engineered heart tissue^69^ .First, iPSC were induced with doxycycline 200 ng/µl for 24 hours to induce the expression of MyoD and Baf60c. The day after, 1.25 × 10^6^ iPSC-committed were resuspended in 77 µl of growth media (SKM02, AMSbiokit) supplemented with hES cell Recovery (Stemgent) and molded in hydrogel composed by 40 µl of Bovine Collagen solution 6mg/ml (Sigma-Aldrich), 17.8 µl of Matrigel Growth Factor reduced (GFR, Corning) 10% v/v 3), 40µl 2X DMEM (Gibco) 4) and 5.2 µl of NaOH 0.05 N). For the generation of MYOrganoids including fibroblasts, 1.25 × 10^5^ (ratio 1:10) fibroblasts were included in the iPSC-committed mix during hydrogel preparation. 180µl of hydrogel was cast into 48-well plate TM5 MyrPlate (Myriamed), containing in each well a pair of flexible poles (static stretchers) that support the growth of the engineered tissue in a ring shape. After 1 hour of polymerization at 37°C, growth media (SKM02, AMSbiokit) was added for 24 hours. On day 2 of the 3D, growth media (SKM02, AMSbiokit) was replaced by differentiation media (SKM03plus, AMSbiokit) and changed every day until day 14. For Condition Media (CM) experiments, single cell-population organoid-like structures were made using the same protocol described above. CM was collected from CTR or DMD fibroblasts-only 3D structure and diluted 1:10 in fresh differentiation media (SKM03plus, AMSbiokit). The diluted media was then used to culture CTR or DMD myotubes-only organoids. CM was collected and used to treat 3D cultures from day 7 until day 13.

### Muscle force analysis

Functional analyses were carried out on day 14 after 3D casting. Contraction experiments were performed using the MyoDynamics Muscle Strip System 840 MD (Danish Myo Technology A/S) and CS4 stimulator (Danish Myo Technology A/S). All functional analysis were performed at 37°C, 5% CO_2_ 95% O_2_, in Tyrode’s solution supplemented with 25 mM NaHCO_3_. Optimal muscle length was determined by gradually stretching the muscle until 1.0 mN of passive tension registered. Functional tests were performed under isometric and eccentric conditions. MYOrganoids were electrically stimulated with 250 pulses of 30V, 4 ms width at the 125 Hz of frequency for both isometric and eccentric contractions. For eccentric analysis, MYOrganoids were 1 mm stretched at the 6.5 mm/s speed during the muscular contraction. Data collection and analysis was done by PowerLab device and LabChart software (ADInstruments, New Zealand) respectively. Each artificial tissue was subjected to 1 isometric contraction, 10 eccentric contractions and 1 isometric contraction. Fatigue is represented as percentage drop force between the first and the last isometric contraction. Where indicated, force is indirectly normalized for the CSA (Cross Section Area) calculated as muscle force (mN) x Lo (mm) x density (mg/mm3)/weight (mg) and expressed as mN/mm2. Engineered muscle density is experimentally determined as 2.089 mg/mm3.

### Immunofluorescence

MYOrganoids that did not undergo functional analysis have been analyzed for immunohistochemistry. Briefly, the artificial tissues were fixed in 4% methanol-free paraformaldehyde (PFA) overnight on day 14. For whole-mount staining, fixed MYOrganoids were permeabilized, stained and cleared with the MACS clearing kit (Miltenyi) according to manufacturer’s instructions. Whole mount-stained organoids are then imaged with confocal microscopy (LEICA STED SP8) at 10X magnification. For staining on transversal or longitudinal sections, fixed MYOrganoids were dehydrated with a gradient of sucrose (7.5%-30%) over day and embedded in OCT matrix in plastic molds. After 24 hours, embedded MYOrganoids were processed with the cryostat (LEICA) with 14 µm thick sections. Slices were then dried and fixed again with 4% methanol-free PFA (Invitrogen). Fixed sections were then blocked with serum cocktail (5% Goat serum and 5% Fetal bovine serum), before being stained overnight at +4°C with primary antibody. After that, slices were washed three times in PBS and hybridized with AlexaFluor secondary antibody according to the host species of the first antibody. Stained slides were then covered with Fluoromont + Dapi (SouthernBiotech) and glass slide 1.5H. For imaging, sections were scanned with AxioScan microscope and confocal Leica SP8. For 2D staining, cells were grown on µ-Dish 35 mm (Ibidi) and then fixed in 4% methanol-free PFA for 7 minutes. For membrane and cytoplasmatic staining, cells were permeabilized with 0.15% Triton X-100 for 10 minutes and then washed with PBS for 5 minutes. For nuclear staining, cells were permeabilized with 0.25% Triton X-100 for 15 minutes. Permeabilized cells were then blocked with serum cocktail (5% Goat serum and 5% Fetal bovine serum), before being stained overnight at +4°C with primary antibody. Cells are then washed three times with PBS and then hybridized with AlexaFluor secondary antibody according to the host species of the first antibody. After three PBS washing, nuclei were stained with Hoechst 33342 (Invitrogen) at the final dilution of 2 µg/ml. For imaging, sections are scanned confocal Leica SP8. All antibodies used for staining are listed in **table S5**.

### Electron Microscopy

Electron microscopy analysis was prospectively performed on MYOtissue specimens that were fixed with glutaraldehyde (2.5%, pH 7.4), post fixed with osmium tetroxide (2%), dehydrated in a graded series of ethanol ranging from 30% to absolute solution and embedded in resin (EMBed-812, Electron Microscopy Sciences, USA). 80 nm thick sections from at least four blocks from CTR iPSC-derived MYOrganoids in presence or absence of CTR fibroblasts were stained with uranyl acetate and lead citrate. The grids were observed using a “JEOL 1400 Flash” electron microscope (120 kV) and were photo documented using a Xarosa camera (Soft Imaging System, France). Images covering whole longitudinal sections were assessed and representative images were used.

### Images analysis

**Myotubes alignment:** FIJI and CellPose were used for image analysis. Myotube alignment was determined by angle measurement and by myotube circularity from cross-section cuts of MYOrganoids. Angles between two myotubes were measured with the “angle tool” function in FIJI, by drawing two lines perpendicular to two adjacent myotubes membrane. Myotubes circularity was determined by custom FIJI script. Briefly, myotubes cross section area was first segmented by pre-trained Cellpose2 cyto2 model ^70^ and then converted into Regions of Interest (ROIs) by Labels_To_Rois.py plugin (Waisman et al. 2021) for subsequent quantification on FIJI (Ferret diameters X and Y). After that, the ratio between Feret’s diameter on axis X and axis Y was assessed. At least 4 frames for 3 biological replicates have been analyzed. **Maturation and fusion index:** Maturation index was assessed by counting nuclei contained in striated myotubes identified by sarcomeric alpha actinin SAA staining in FIJI software and normalized for the total number of nuclei contained in myotubes. Fusion index has been calculated counting myotubes myosin heavy chain (MyHC) positive containing 1, 2, 3 or more than 3 nuclei as a percentage of the total number of nuclei. At least 4 frames for 3 biological replicates have been analyzed. **Z-disk length analysis:** Z-disk length has been assessed by drawing straight lines in FIJI and by measuring the length in at least 10 mature myotubes per condition. **Phospho-SMAD3 quantification:** Phospho-SMAD3 (pSMAD3) has been measured as a percentage of positive nuclei of myosin heavy chain positive cells (MyHC+, i.e. myotubes) or myosin heavy chain negative cells (MyHC-, i.e. fibroblasts). At least 4 frames for 3 biological replicates have been analyzed. **Dystrophin, α-Dystroglycan and β-Dystroglycan quantification:** MYOtissue cross sections were stained sarcomeric alpha-actinin (SAA) for myotubes cytosol labeling. The Cellpose2 cyto2 model 10 was fine-tuned on manually myofiber-labeled images based on SAA staining (hyperparameters: n_epochs=200, learning_rate=0.05, weight_decay=0.0001). The labeled dataset used in fine-tuning was prepared in such a way that the model can simultaneously segment myotubes and ignore low-quality staining areas. Fine-tuned models were then used to extract myotubes masks. Reconstruction of myotubes masks was done using the cellpose package (Stringer et al. 2021). Reconstructed masks were then converted into Regions of Interest (ROIs) for subsequent quantification (each ROI corresponds to an individual myofiber) using the Labels_To_Rois.py FIJI plugin ^71^. The generated ROIs were used for subsequent quantification using FIJI macro. Positive ROIs for dystrophin / alpha-dystroglycan / beta-dystroglycan signal were counted and represented as a percentage of the total ROIs of one image.

### Gene expression analysis

For gene expression analysis, bulk RNA was isolated from MYOrganoids by RNeasy micro kit (QIAGEN) according to manufacturer’s instructions, controlled and quantified by Nanodrop. Around 0.5-1µg of RNA was retro-transcribed to cDNA thanks to the RevertAid H Minus First Strand cDNA Synthesis Kit (Invitrogen). Droplet digital PCR was performed to assess the expression of myogenic factors (MYOGENIN, MYH2, MYH7, MCK), of fibrotic markers (COL1A1, FN1, TGF-β1) and µDystrophin, thanks to the QX200™ ddPCR™ EvaGreen Supermix (Biorad). Gene expression results in copy/µl (cp/µl) are represented as fold change to the expression of GAPDH gene. Primers used are listed in **table S6**.

### ELISA assay

Media was collected before muscle functional analysis after being in culture for 24 hours. Secreted human Fibronectin 1 (FN1), TGF-β and Collagen IV were assessed by ELISA kit (**table S7**) from MYOrganoids supernatant.

### AAV production and MYOrganoids infection

Recombinant AAVs were produced as previously described (Bourg et al. 2022) using AAV9 serotype. Purification was performed using affinity chromatography and titration was done by ddPCR using transgene-specific primers. For optimization of infection, an AAV9-CMV-GFP construct was used. The micro-dystrophin transgene used in the study, under the control of spc512 promoter, was an optimized version of the construct used for GENETHON’s preclinical investigation and clinical trial ^17^, with deletion from spectrin-like repeats 4 to 23 and full C-terminal truncation, here referred as µDys. Infection in MYOrganoids was performed by delivering the AAV9 particles diluted into the differentiation media at day 7, at two different doses: 1E+9 vg/MYOtissue (low dose) and 5E+10 vg/MYOtissue (high dose). Media was replaced after 24 hours from the infection and changed daily until day 14.

### Viral copy number analysis

Viral DNA was extracted from mature MYOrganoids by NucleoMag Pathogen kit (Macherey Nagel) using Kingfisher instrument (Thermofisher). DNA yield and purity was assessed by Nanodrop; viral copy number (VCN) was identified by droplet digital PCR using supermix for probe (Biorad). Results are shown as copy number variation using P0 as reference DNA. Primers used are listed in **table S6**.

### Capillary western blot analysis

MYOrganoids proteins were extracted in RIPA buffer supplemented with Protease Inhibitor Cocktail EDTA-free (Roche) and Benzonase by homogenization. Total proteins were then quantified by BCA method, thanks to the Pierce 660 protein assay kit (Invitrogen) according to manufacturer’s instructions. Protein detection has been performed by capillary western blot, thanks to the JESS protein simple (Bio-techne), according to manufacturer’s directions. Dystrophin detection (both full-length and µDystrophin) has been performed by the antibody DysB (**table S5**) and its expression has been quantified by total protein normalization.

### RNA sequencing and transcriptomic analysis

The RNA quality of samples was verified using the Bioanalyzer 2100 (Agilent) and Qubit fluorometric quantification (ThermoFisher Scientific). The samples that had an RNA integrity number higher than 9 were used for RNA sequencing (Genewiz). The Stranded Total RNA Library Prep Kit (Illumina) was used to create sequencing libraries, which were sequenced following the Illumina protocol on the NovaSeq instrument (Illumina), resulting in approximately 20 million paired-end reads per library. The paired-end reads were filtered and subjected to quality control using fastp (Chen et al. 2018). They were then mapped to the GRCh38/hg38 genome using HISAT2 ^72^ count tables were generated using htseq-count ^73^. Differentially expressed genes (DEGs) were identified using the DESeq2 R package with p value adjusted by Benjamin-Hochberg procedure less than 0.05. Pathway analysis was carried out in R-Studio (version 4.0.3) using functional class scoring with Gene Set Enrichment Analysis ^74,75^.

### Statistical analysis

All data were analyzed by GraphPad Prism v10.2.2 software. Parametric tests such as t-tests and ANOVA were used for statistical comparison. To compare two groups, unpaired one-tail t-test was used; to compare multiple groups, we used one-way ANOVA with Tukey’s correction for multiple comparison tests, with the assumption of Gaussian distribution of residuals. Results were considered significantly different at p < 0.05. Graphs were generated using Graphpad Prism v10.2.2 or R version 4.0.3, The figures display the mean ± standard error of the mean (SEM) as described in the figure legend.

