## Supplementary info for "Disease exacerbation by fibroblast inclusion in Duchenne Muscular Dystrophy MYOrganoids reveals limitations of microdystrophin therapeutic efficacy"

#### Supplementary Materials

**This PDF file includes:**

**Supplementary figures 1-8**

**Supplementary table 1** List of GO biological processes differentially expressed between DMD organoids (DMDdEx8-9) and isogenic control (IsoCTR)

**Supplementary table 2** List of significant (p.adj < 0.05) differentially expressed genes (DEGs) between DMD organoids treated with µDystrophin (dEx8-9-µDys) and untreated (dEx8-9)

**Supplementary table 3** List of GO biological processes differentially expressed between DMD organoids treated with µDystrophin (dEx8-9-µDys) and isogenic control (IsoCTR)

**Supplementary table 4** Cell lines list

**Supplementary table 5** Antibodies used for immunofluorescence and capillary western blot

**Supplementary table 6** List of primers used for viral copy number analysis and gene expression

**Supplementary table 7** ELISA kit used for secreted protein analysis

### Supplementary Figure 1

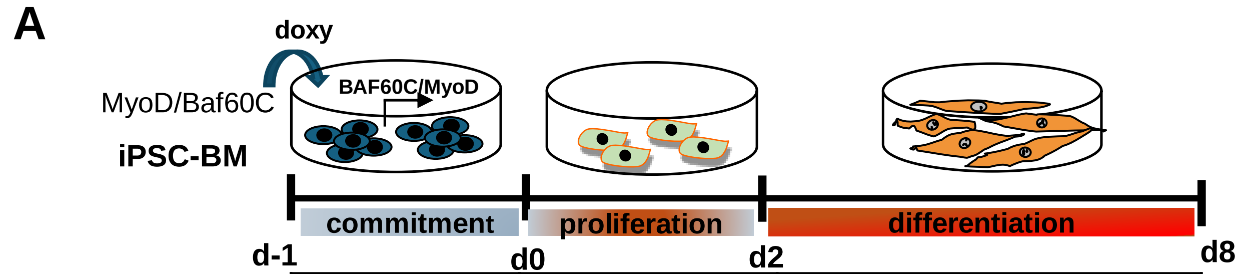


| | | |
|---|---|---|
| iPSC medium | GM | DM |
| iPSC medium | GM | SK03plus medium |
| iPSC medium | SK02 medium | SK03plus medium |

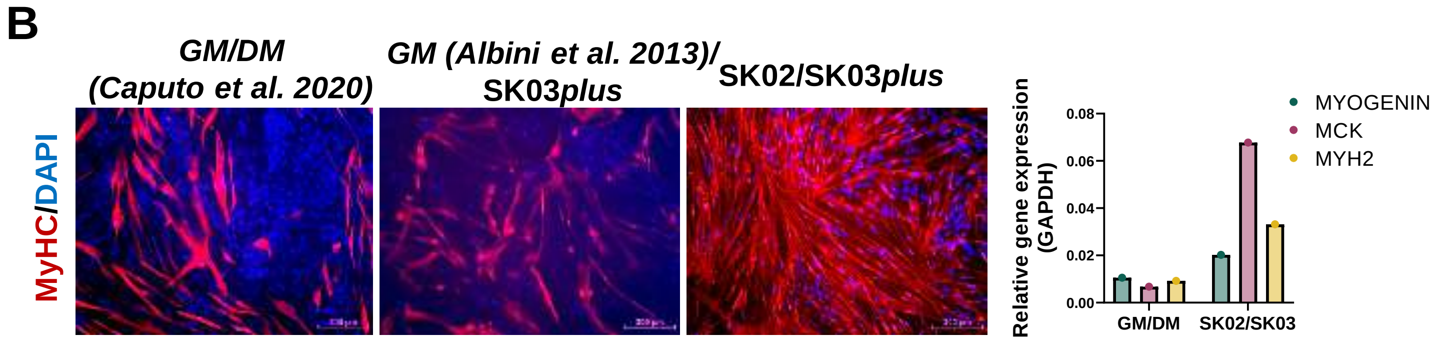


**Supplementary figure 1. Optimization of direct myogenic differentiation of iPSC (A)** Scheme of the protocol for 2D direct muscle differentiation from iPSC, adapted from Caputo et al. 2020. GM: homemade growth medium; DM: homemade differentiation medium. Commercial proliferation (SK02) and differentiation (SK03+) media were used to achieve optimal differentiation. (**B**) Myogenic differentiation evaluated by MyHC immunostaining, whose representative images are shown, and by RT-ddPCR analysis of early (*MYOGENIN*) and late (*MCK, Muscle Creatine-kinase; MYH2, Myosin Heavy Chain 2*) myogenic markers.

### Supplementary Figure 2

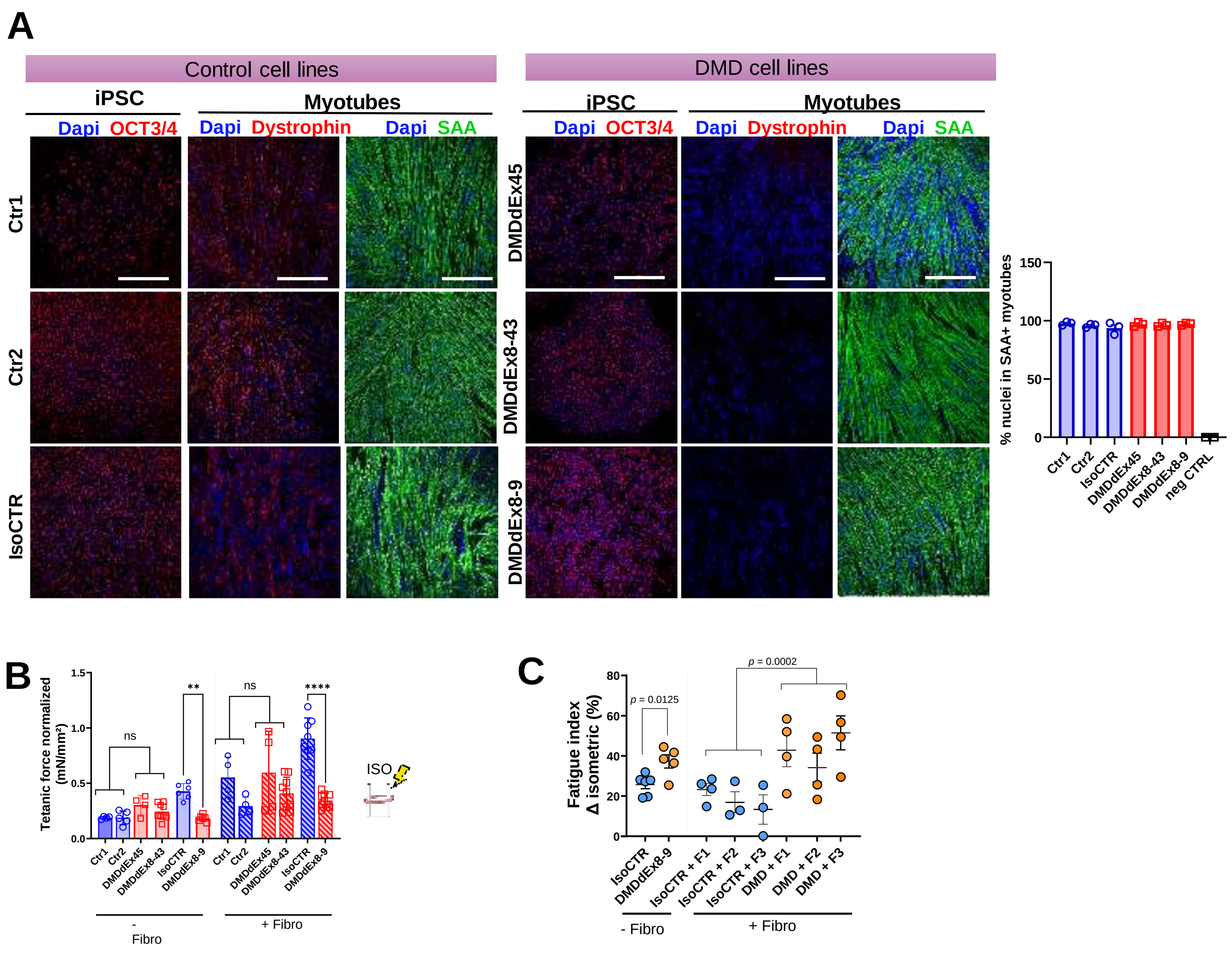


**Supplementary figure 2. Myotubes differentiation assessment in monolayer culture and characterization of MYOrganoids including and not including fibroblasts (A)** Oct3/4, dystrophin and sarcomeric alpha-actinin (SAA) staining in hIPSC (OCT3/4) and in myotubes (dystrophin and SAA) in Ctr1, Ctr2, DMDdEx45, DMDdEx8-43 and the isogenic iPSC DMDdEx8-9 with their corrected control (IsoCTR). Scale bar: 150 µm. Maturation index has been calculated as a percentage of nuclei in SAA-positive myotubes. (**B**) Isometric tetanic force of Ctr and DMD MYOrganoids with and without fibroblasts showing no difference upon a single isometric pulse. Results are normalized for cross-section area. (**C)** Fatigue index, in organoids generated from IsoCTR and DMDdEx8-9-derived muscle cells without fibro (- Fibro) and with 3 Ctr and 3 DMD fibroblasts obtained from different genetic sources (F1+F2+F3). N=3-6. Data are presented as means +/- SEM. Statistical analysis was performed with an ordinary one-way ANOVA test (*p ≤0.05, **p ≤ 0.01, ***p ≤ 0.001, ****p ≤ 0.0001, ns = not significant) with multiple comparisons corrected with Tukey's test.

### Supplementary Figure 3

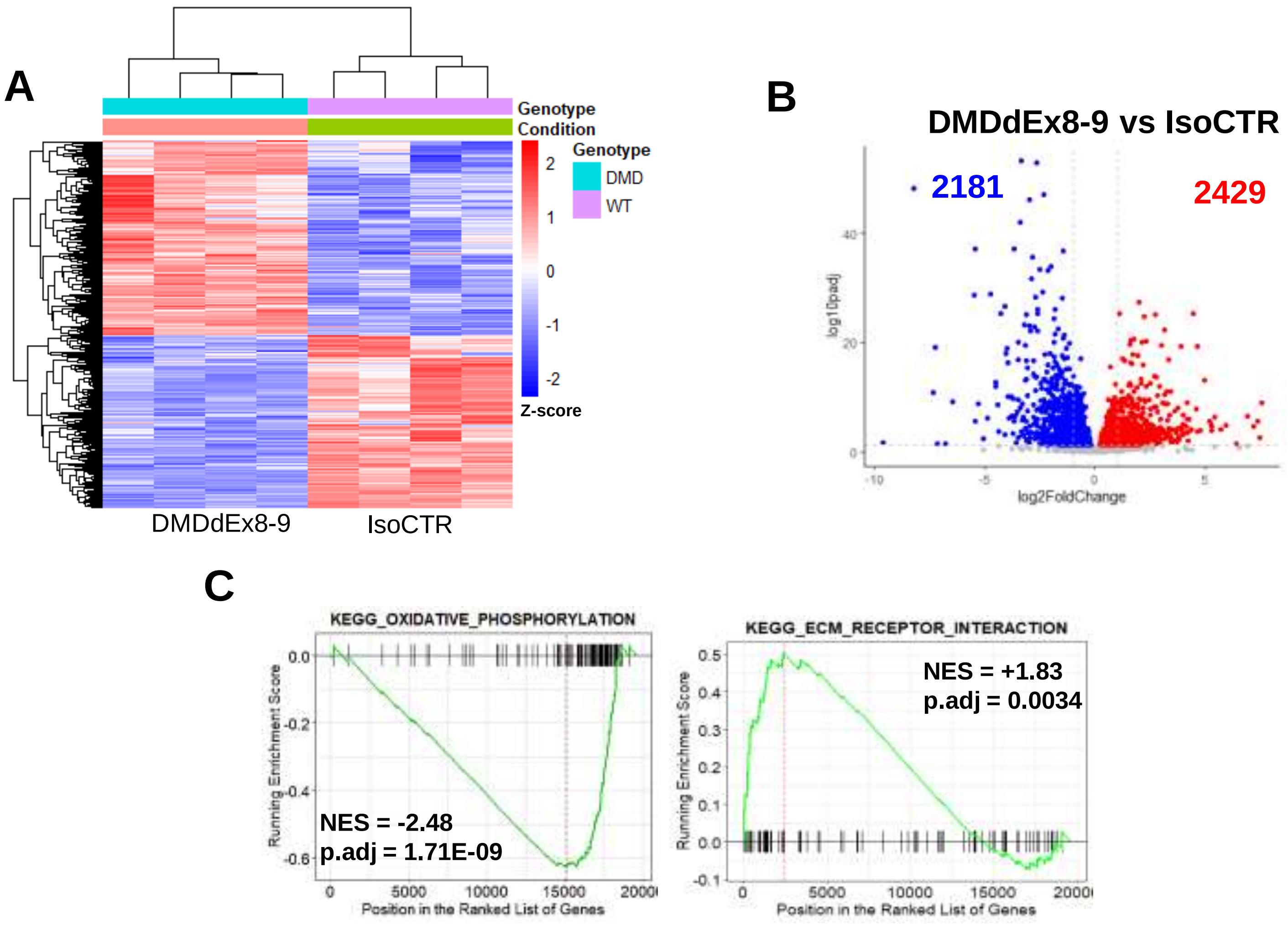


**Supplementary figure 3. Transcriptomic characterization of DMD MYOrganoids. (A**) Clustered heatmap of significant differentially expressed genes (DEGs) between DMDdEx8-9 and isogenic control (IsoCTR) MYOrganoid (**B**) Vulcano plot representing DEGs between DMDdEx8-9 and IsoCTR MYOrganoids (**C)** Kyoto encyclopedia of genes and genome (KEGG) enrichment analysis of DEGs between DMDdEx8-9 and IsoCTR MYOrganoids.

### Supplementary Figure 4

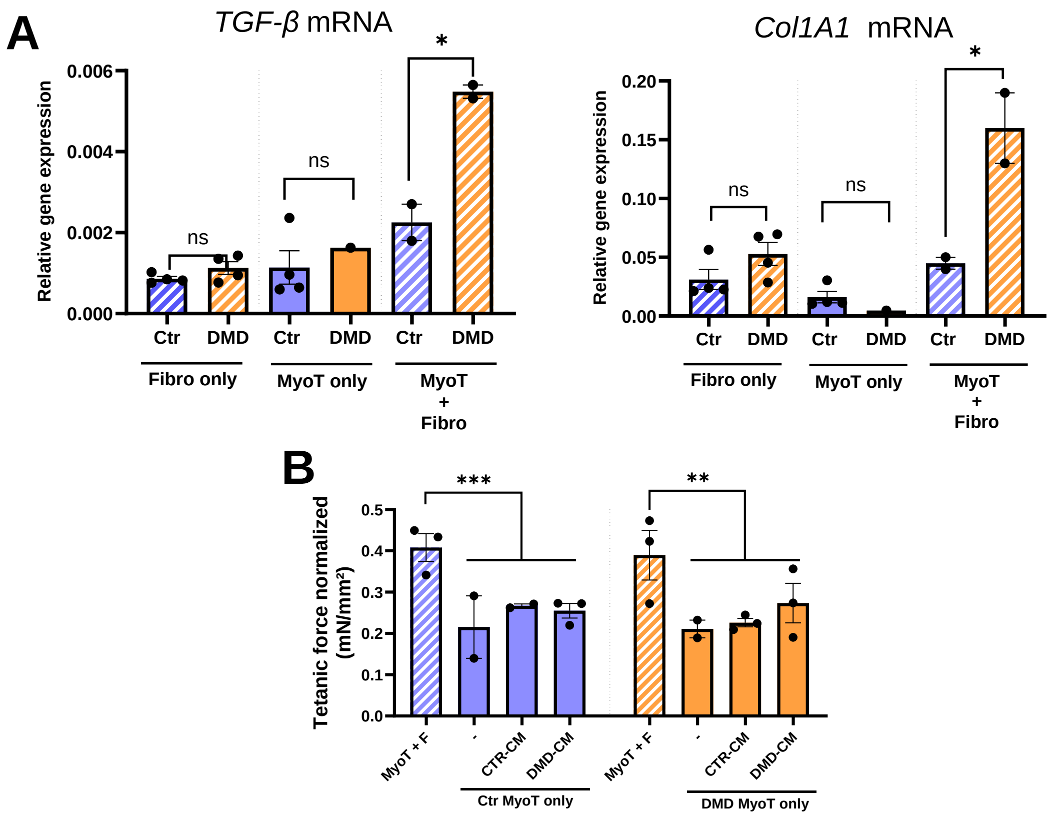


**Supplementary figure 4. Fibroblast role in the modeling of DMD phenotype (A**) Gene expression analysis of TGF-β and Collagen 1A1 (Col1A1) in fibroblasts only (Fibro-only), myotubes-only (MyoT-only) 3D cell cultures and MYOrganoids containing myotubes with fibroblasts (MyoT+Fibro) in control (Ctr) and DMD conditions (**B**) Isometric tetanic force normalized for cross-section area of MYOrganoids (MyoT + Fibro), and MyoT only 3D cell culture untreated (-) and treated with control conditioned media from control fibro-only tissues (CTR-CM) or DMD fibro-only (DMD-CM).

### Supplementary Figure 5

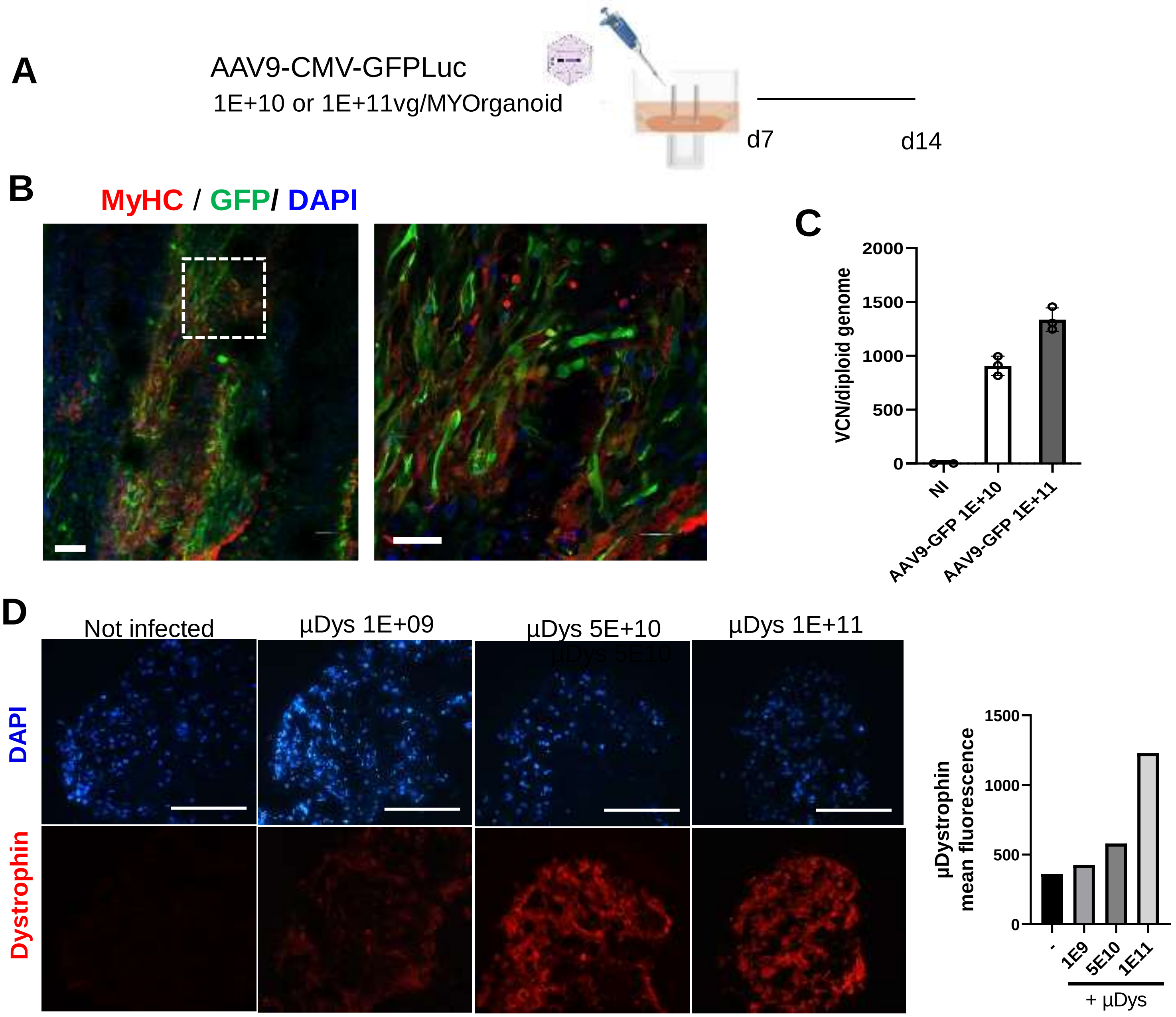


**Supplementary figure 5. Optimization of AAV infection in MYOrganoids. (A)** Infection method using AAV9-CMV-GFPluc. (**B**) Representative images showing infection efficiency using whole-mount staining with GFP antibody. (**C**) VCN quantification from MYOrganoids infected with AAV9-CMV-GFP-Luc at two different doses: 1^10 and 1^11 viral genome per organoid. (**D**) Myofiber transduction, evaluated by dystrophin staining, following increasing dosing of AAV9-Sp512-µDys vector. DAPI stains nuclei. Scale bar: 100 µm.

### Supplementary Figure 6

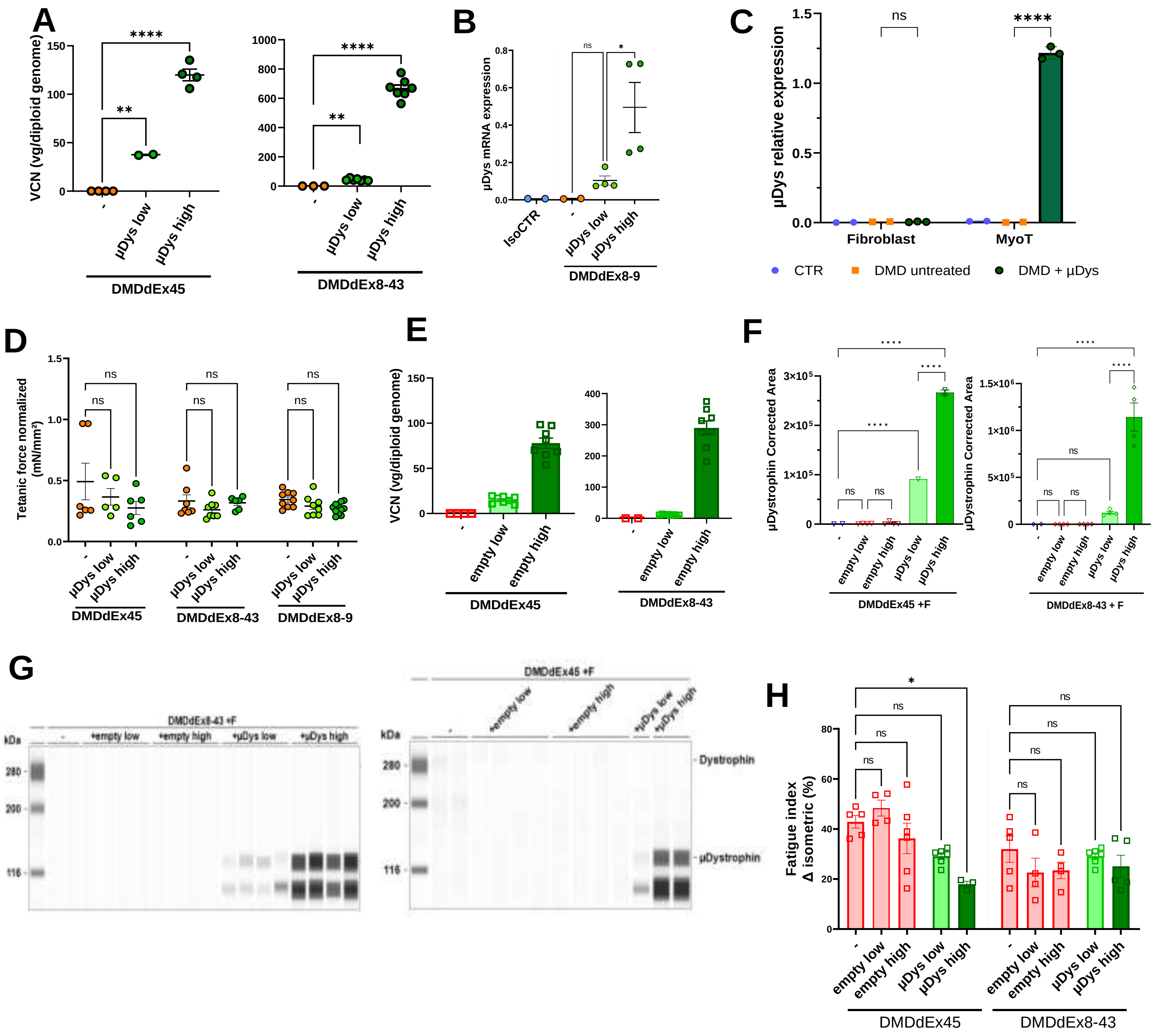


**Supplementary figure 6. Efficacy of µDystrophin gene transfer and effect of empty vector on MYOrganoids functionality. (A)** Viral copy number (VCN) of µDys in DMDdEx8-43 and DMDdEx45 MYOrganoids infected with AAV9-µDys indicated as viral genome (vg) per diploid genome (**B**) relative mRNA expression of transgene in DMD dEx8-9 organoids upon µDys. (**C**) µDystrophin protein expression evaluated by Capillary western blot in fibroblasts or iPSC-derived muscle cells, infected or not with AAV9-µDys (D) (**D**) Tetanic force normalized in DMDdEx8-9 DMDdEx8-43 and DMDdEx45 MYOrganoids infected with AAV9-µDys, subjected to a first isometric contraction. (**E**) VCN analysis of empty vector genome in DMDdEx8-43 and DMDdEx45 MYOtissues infected with AAV9-empty vector (**F**) Capillary western blot quantification of µDystrophin in DMDdEx45 MYOrganoids or DMDdEx8-43 MYOrganoids infected at two doses with AAV9-µDys and AAV9-empty vector **(G)** Run pictures of capillary western blot of DMDdEx45 MYOrganoids or DMDdEx8-43 MYOrganoids infected at two doses with AAV9-µDys and AAV9-empty vector (**H**) Fatigue index in DMDdEx45 and DMDdEx8-43 MYOtissues infected at two doses with an empty vector and with µDystrophin. For all panels, data are presented as means +/- SEM. Statistical analysis was performed with an ordinary one-way ANOVA test (*p ≤0.05, **p ≤ 0.01, ***p ≤ 0.001, ****p ≤ 0.0001, ns = not significant).

### Supplementary Figure 7

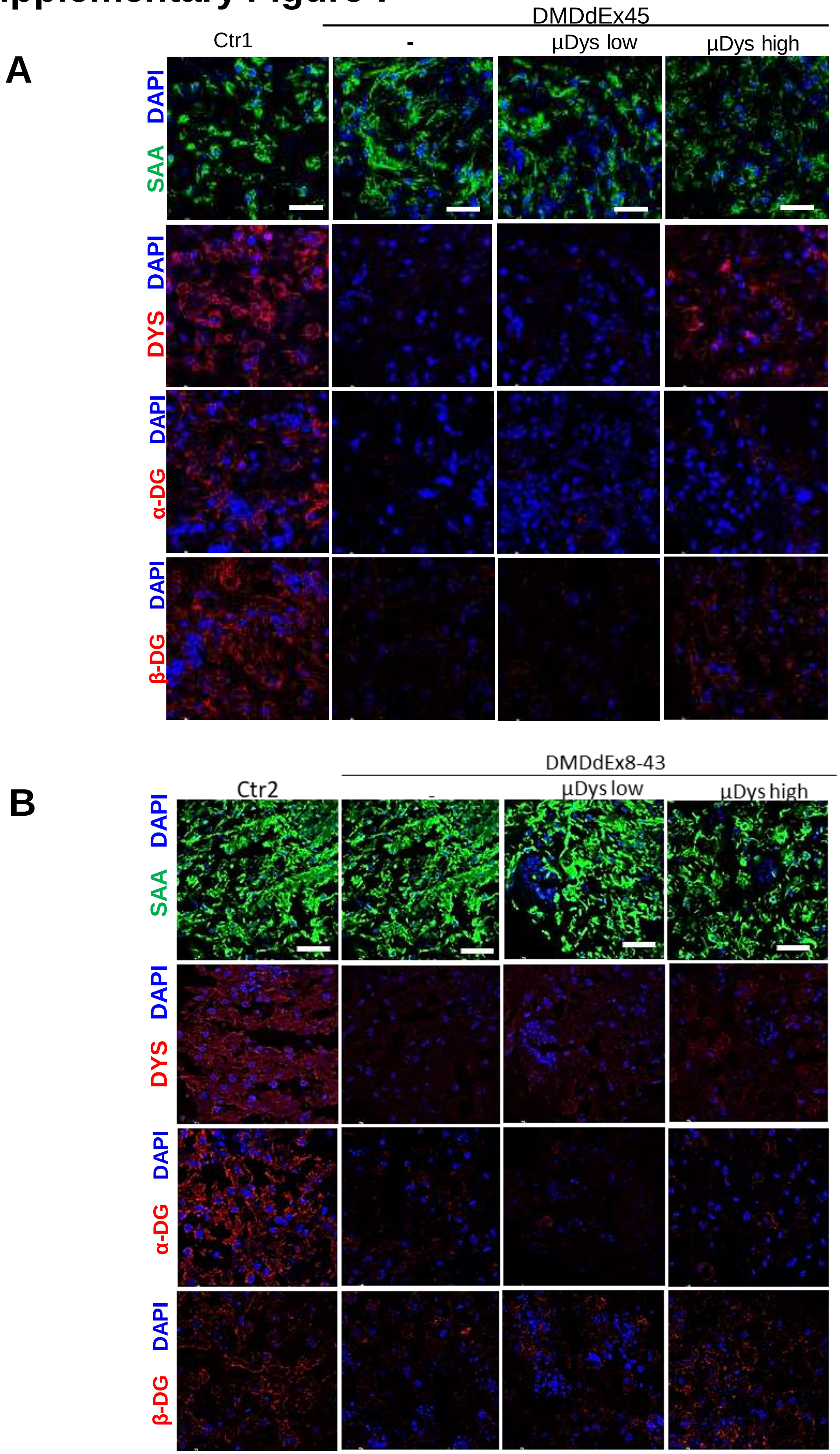


**Supplementary figure 7. Efficacy of µDystrophin infection and restoration of dystrophin Glycoprotein complex in additional iPSC derived-MYOrganoids. (A)** Sarcomeric alpha-actinin (SAA), dystrophin (dys), α-dystroglycan (α-DG) and β-dystroglycan (β-DG) immunostaining in Ctr1, DMDdEx45 untreated and infected at two doses (low and high) of µDystrophin (µDys). Scale bar: 50 µm. Relative quantifications have been performed by calculating the percentage of DYS/a-DG/b-DG and SAA positive myotubes **(B)** Sarcomeric alpha-actinin (SAA), dystrophin (dys), α-dystroglycan (α-DG) and β-dystroglycan (β-DG) immunostaining in Ctr2, and DMDdEx8-43 untreated and infected at low and high doses of µDys ). Scale bar: 50 µm.

### Supplementary Figure 8

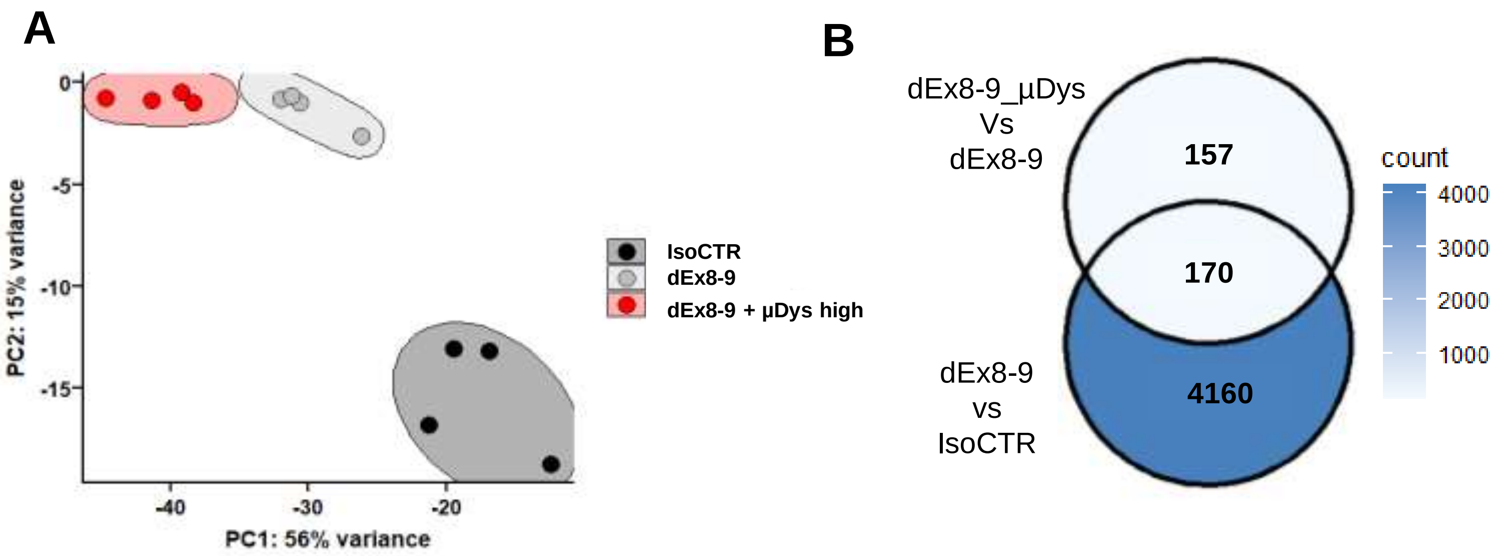


**Supplementary figure 8. Transcriptomic analysis in DMDdEx8-9 treated with µDys and comparison with untreated and isogenic control (A)** Principal component analysis (PCA) of DMDdEx8-9 MYOrganoids (dEx8-9) treated with µDys (dEx8-9 + µDys high) compared with isogenic control (IsoCTR) **(B)** Venn diagram of differentially expressed genes (DEGs) between dEx8-9 MYOrganoids treated with µDys and untreated (dEx8-9_µDys vs dEx8-9) and dEx8-9 MYOrganoids vs isogenic control (dEx8-9 vs IsoCTR).

**Supplementary table 1: List of GO biological processes differentially expressed between DMD organoids (DMDdEx8-9) and isogenic control (IsoCTR)**

| Description | setSize | NES | p.adjust | qvalue |
|---|---|---|---|---|
| GOBP_MUSCLE_SYSTEM_PROCESS | 390 | -2,103 | 1,31E-09 | 1,22E-09 |
| GOBP_MUSCLE_CONTRACTION | 308 | -2,170 | 2,18E-09 | 2,03E-09 |
| GOBP_NEGATIVE_REGULATION_OF_MEGAKARYOCYTE_DIFFERENTIATION | 17 | 2,429 | 1,92E-08 | 1,79E-08 |
| GOBP_CYTOPLASMIC_TRANSLATION | 146 | -2,375 | 3,62E-08 | 3,38E-08 |
| GOBP_MITOCHONDRIAL_RESPIRATORY_CHAIN_COMPLEX_ASSEMBLY | 89 | -2,454 | 2,86E-07 | 2,66E-07 |
| GOBP_AEROBIC_RESPIRATION | 166 | -2,247 | 2,86E-07 | 2,66E-07 |
| GOBP_DNA_REPLICATION_DEPENDENT_CHROMATIN_ORGANIZATION | 32 | 2,473 | 8,37E-07 | 7,81E-07 |
| GOBP_CELLULAR_RESPIRATION | 208 | -2,085 | 3,31E-06 | 3,09E-06 |
| GOBP_STRIATED_MUSCLE_CONTRACTION | 161 | -2,132 | 5,59E-06 | 5,21E-06 |
| GOBP_OXIDATIVE_PHOSPHORYLATION | 119 | -2,216 | 6,40E-06 | 5,97E-06 |
| GOBP_DNA_CONFORMATION_CHANGE | 307 | 1,867 | 1,28E-05 | 1,19E-05 |
| GOBP_NADH_DEHYDROGENASE_COMPLEX_ASSEMBLY | 53 | -2,451 | 1,33E-05 | 1,24E-05 |
| GOBP_ENERGY_DERIVATION_BY_OXIDATION_OF_ORGANIC_COMPOUNDS | 281 | -1,946 | 1,33E-05 | 1,24E-05 |
| GOBP_SKELETAL_MUSCLE_CONTRACTION | 41 | -2,383 | 3,21E-05 | 2,99E-05 |
| GOBP_STRIATED_MUSCLE_CELL_DEVELOPMENT | 64 | -2,305 | 1,05E-04 | 9,82E-05 |
| GOBP_REGULATION_OF_MUSCLE_CONTRACTION | 143 | -2,058 | 1,16E-04 | 1,08E-04 |
| GOBP_REGULATION_OF_MUSCLE_SYSTEM_PROCESS | 209 | -1,926 | 1,38E-04 | 1,29E-04 |
| GOBP_REGULATION_OF_SYSTEM_PROCESS | 482 | -1,671 | 1,38E-04 | 1,29E-04 |
| GOBP_ATP_METABOLIC_PROCESS | 240 | -1,868 | 1,76E-04 | 1,64E-04 |
| GOBP_MITOCHONDRIAL_ELECTRON_TRANSPORT_NADH_TO_UBIQUINONE | 43 | -2,292 | 2,94E-04 | 2,74E-04 |
| GOBP_DNA_REPLICATION_INDEPENDENT_CHROMATIN_ORGANIZATION | 34 | 2,221 | 2,94E-04 | 2,74E-04 |

| | | | | |
|---|---|---|---|---|
| GOBP_DETECTION_OF_STIMULUS_INVOLVED_IN_SENSORY_PERCEPTION | 118 | 2,017 | 2,94E-04 | 2,74E-04 |
| GOBP_MITOTIC_NUCLEAR_DIVISION | 282 | 1,814 | 2,94E-04 | 2,74E-04 |
| GOBP_MULTICELLULAR_ORGANISMAL_MOVEMENT | 53 | -2,297 | 3,32E-04 | 3,10E-04 |
| GOBP_CHROMOSOME_SEGREGATION | 315 | 1,731 | 3,65E-04 | 3,41E-04 |
| GOBP_SARCOMERE_ORGANIZATION | 40 | -2,259 | 6,36E-04 | 5,94E-04 |
| GOBP_ATP_SYNTHESIS_COUPLED_ELECTRON_TRANSPORT | 77 | -2,149 | 6,36E-04 | 5,94E-04 |
| GOBP_HEART_PROCESS | 224 | -1,807 | 6,36E-04 | 5,94E-04 |
| GOBP_NUCLEAR_CHROMOSOME_SEGREGATION | 257 | 1,767 | 6,36E-04 | 5,94E-04 |
| GOBP_DNA_DEPENDENT_DNA_REPLICATION | 156 | 1,843 | 6,98E-04 | 6,51E-04 |
| GOBP_RESPIRATORY_ELECTRON_TRANSPORT_CHAIN | 98 | -2,053 | 7,47E-04 | 6,97E-04 |
| GOBP_SISTER_CHROMATID_SEGREGATION | 191 | 1,862 | 9,30E-04 | 8,67E-04 |
| GOBP_ACTIVATION_OF_GTPASE_ACTIVITY | 107 | 1,980 | 1,18E-03 | 1,10E-03 |
| GOBP_REGULATION_OF_MEGAKARYOCYTE_DIFFERENTIATION | 35 | 2,169 | 1,21E-03 | 1,13E-03 |
| GOBP_REGULATION_OF_HEART_CONTRACTION | 181 | -1,814 | 1,30E-03 | 1,21E-03 |
| GOBP_SARCOPLASMIC_RETICULUM_CALCIUM_ION_TRANSPORT | 36 | -2,192 | 1,38E-03 | 1,29E-03 |
| GOBP_MITOTIC_SISTER_CHROMATID_SEGREGATION | 166 | 1,857 | 1,39E-03 | 1,30E-03 |
| GOBP_REGULATION_OF_BLOOD_CIRCULATION | 219 | -1,775 | 1,61E-03 | 1,50E-03 |
| GOBP_RIBOSOME_BIOGENESIS | 298 | -1,676 | 1,82E-03 | 1,70E-03 |
| GOBP_REGULATION_OF_STRIATED_MUSCLE_CONTRACTION | 83 | -2,048 | 1,89E-03 | 1,76E-03 |
| GOBP_MITOCHONDRIAL_TRANSLATION | 72 | -2,074 | 2,04E-03 | 1,90E-03 |
| GOBP_SPINDLE_ORGANIZATION | 199 | 1,772 | 2,06E-03 | 1,92E-03 |
| GOBP_DNA_PACKAGING | 212 | 1,765 | 2,06E-03 | 1,92E-03 |
| GOBP_ORGANELLE_FISSION | 447 | 1,568 | 2,50E-03 | 2,33E-03 |
| GOBP_COLLAGEN_FIBRIL_ORGANIZATION | 60 | 2,023 | 2,67E-03 | 2,49E-03 |

| | | | | |
|---|---|---|---|---|
| GOBP_MITOCHONDRION_ORGANIZATION | 492 | -1,519 | 4,90E-03 | 4,57E-03 |
| GOBP_STRIATED_MUSCLE_CELL_DIFFERENTIATION | 251 | -1,659 | 5,62E-03 | 5,24E-03 |
| GOBP_EXTERNAL_ENCAPSULATING_STRUCTURE_ORGANIZATION | 281 | 1,665 | 5,69E-03 | 5,31E-03 |
| GOBP_GENERATION_OF_PRECURSOR_METABOLITES_AND_ENERGY | 436 | -1,524 | 5,69E-03 | 5,31E-03 |
| GOBP_CELLULAR_COMPONENT_ASSEMBLY_INVOLVED_IN_MORPHOGENESIS | 99 | -1,941 | 6,78E-03 | 6,33E-03 |
| GOBP_ACTIN_MYOSIN_FILAMENT_SLIDING | 14 | -2,098 | 7,64E-03 | 7,13E-03 |
| GOBP_MUSCLE_CELL_DEVELOPMENT | 168 | -1,754 | 1,07E-02 | 9,94E-03 |
| GOBP_DNA_GEOMETRIC_CHANGE | 94 | 1,853 | 1,15E-02 | 1,08E-02 |
| GOBP_REGULATION_OF_CATION_TRANSMEMBRANE_TRANSPORT | 298 | -1,571 | 1,20E-02 | 1,12E-02 |
| GOBP_VENTRICULAR_CARDIAC_MUSCLE_TISSUE_DEVELOPMENT | 52 | -1,998 | 1,23E-02 | 1,15E-02 |
| GOBP_SPINDLE_ASSEMBLY | 134 | 1,760 | 1,23E-02 | 1,15E-02 |
| GOBP_INTERLEUKIN_1_MEDIATED_SIGNALING_PATHWAY | 26 | -2,113 | 1,31E-02 | 1,22E-02 |
| GOBP_THYROID_HORMONE_METABOLIC_PROCESS | 21 | 2,007 | 1,41E-02 | 1,31E-02 |
| GOBP_NEUROMUSCULAR_PROCESS | 138 | -1,755 | 1,46E-02 | 1,36E-02 |
| GOBP_DNA_REPLICATION | 275 | 1,618 | 1,65E-02 | 1,54E-02 |
| GOBP_REGULATION_OF_CALCIUM_ION_IMPORT | 36 | -1,988 | 1,66E-02 | 1,55E-02 |
| GOBP_DETECTION_OF_STIMULUS | 213 | 1,629 | 1,66E-02 | 1,55E-02 |
| GOBP_CELL_FATE_COMMITMENT | 215 | 1,628 | 1,80E-02 | 1,68E-02 |
| GOBP_RIBOSOME_ASSEMBLY | 59 | -1,943 | 1,87E-02 | 1,74E-02 |
| GOBP_CARDIAC_VENTRICLE_MORPHOGENESIS | 63 | -1,935 | 1,87E-02 | 1,75E-02 |
| GOBP_CATECHOLAMINE_SECRETION | 47 | -1,934 | 1,87E-02 | 1,75E-02 |
| GOBP_MITOCHONDRIAL_GENE_EXPRESSION | 104 | -1,795 | 1,92E-02 | 1,79E-02 |
| GOBP_POSITIVE_REGULATION_OF_CELL_CYCLE_PROCESS | 211 | 1,637 | 1,92E-02 | 1,79E-02 |
| GOBP_RIBONUCLEOPROTEIN_COMPLEX_BIOGENESIS | 422 | -1,472 | 1,92E-02 | 1,79E-02 |

| | | | | |
|---|---|---|---|---|
| GOBP_SISTER_CHROMATID_COHESION | 53 | 1,919 | 1,96E-02 | 1,83E-02 |
| GOBP_ATP_BIOSYNTHETIC_PROCESS | 48 | -1,903 | 2,27E-02 | 2,12E-02 |
| GOBP_BIOMINERALIZATION | 143 | 1,724 | 2,27E-02 | 2,12E-02 |
| GOBP_RECOMBINATIONAL_REPAIR | 156 | 1,630 | 2,27E-02 | 2,12E-02 |
| GOBP_POSITIVE_REGULATION_OF_CELL_CYCLE | 285 | 1,543 | 2,27E-02 | 2,12E-02 |
| GOBP_SENSORY_PERCEPTION_OF_SMELL | 68 | 1,868 | 2,43E-02 | 2,26E-02 |
| GOBP_REGULATION_OF_PROTEIN_DEPOLYMERIZATION | 85 | -1,844 | 2,44E-02 | 2,28E-02 |
| GOBP_MEGAKARYOCYTE_DIFFERENTIATION | 55 | 1,891 | 2,47E-02 | 2,30E-02 |
| GOBP_MUSCLE_ADAPTATION | 98 | -1,795 | 2,47E-02 | 2,30E-02 |
| GOBP_MORPHOGENESIS_OF_AN_EPITHELIAL_FOLD | 20 | 1,953 | 2,47E-02 | 2,30E-02 |
| GOBP_REGULATION_OF_SKELETAL_MUSCLE_CONTRACTION | 16 | -2,009 | 2,64E-02 | 2,46E-02 |
| GOBP_ATP_SYNTHESIS_COUPLED_PROTON_TRANSPORT | 22 | -2,005 | 2,77E-02 | 2,58E-02 |
| GOBP_ACUTE_INFLAMMATORY_RESPONSE | 78 | -1,847 | 2,77E-02 | 2,58E-02 |
| GOBP_ACTIN_MEDIATED_CELL_CONTRACTION | 94 | -1,774 | 2,85E-02 | 2,66E-02 |
| GOBP_REGULATION_OF_ATP_DEPENDENT_ACTIVITY | 70 | -1,855 | 2,89E-02 | 2,69E-02 |
| GOBP_CARDIAC_MUSCLE_CONTRACTION | 123 | -1,751 | 2,89E-02 | 2,69E-02 |
| GOBP_RIBOSOMAL_LARGE_SUBUNIT_BIOGENESIS | 71 | -1,905 | 2,95E-02 | 2,75E-02 |
| GOBP_AMINE_TRANSPORT | 80 | -1,825 | 2,95E-02 | 2,75E-02 |
| GOBP_NEGATIVE_REGULATION_OF_PROTEIN_CONTAINING_COMPLEX_DISASSEMBLY | 79 | -1,799 | 2,95E-02 | 2,75E-02 |
| GOBP_CARDIAC_VENTRICLE_DEVELOPMENT | 115 | -1,698 | 2,95E-02 | 2,75E-02 |
| GOBP_MICROTUBULE_CYTOSKELETON_ORGANIZATION_INVOLVED_IN_MITOSIS | 150 | 1,691 | 3,09E-02 | 2,88E-02 |
| GOBP_PROTEIN_DNA_COMPLEX_ASSEMBLY | 172 | 1,643 | 3,14E-02 | 2,93E-02 |
| GOBP_MUSCLE_CELL_DIFFERENTIATION | 345 | -1,497 | 3,14E-02 | 2,93E-02 |
| GOBP_REGULATION_OF_EMBRYONIC_DEVELOPMENT | 60 | 1,818 | 3,37E-02 | 3,15E-02 |

| | | | | |
|---|---|---|---|---|
| GOBP_PROTON_TRANSMEMBRANE_TRANSPORT | 123 | -1,728 | 3,37E-02 | 3,15E-02 |
| GOBP_TELOMERE_ORGANIZATION | 160 | 1,649 | 3,37E-02 | 3,15E-02 |
| GOBP_NEGATIVE_REGULATION_OF_ACTIN_FILAMENT_DEPOLYMERIZATION | 43 | -1,852 | 3,40E-02 | 3,17E-02 |
| GOBP_CELLULAR_RESPONSE_TO_INTERLEUKIN_1 | 86 | -1,763 | 3,40E-02 | 3,17E-02 |
| GOBP_HOMOPHILIC_CELL_ADHESION_VIA_PLASMA_MEMBRANE_ADHESION_MOLECULES | 157 | 1,654 | 3,45E-02 | 3,22E-02 |
| GOBP_POSITIVE_REGULATION_OF_CALCIUM_ION_IMPORT | 18 | -1,939 | 3,65E-02 | 3,40E-02 |
| GOBP_VENTRICULAR_CARDIAC_MUSCLE_TISSUE_MORPHOGENESIS | 44 | -1,916 | 4,04E-02 | 3,77E-02 |
| GOBP_MITOCHONDRIAL_ATP_SYNTHESIS_COUPLED_PROTON_TRANSPORT | 16 | -1,961 | 4,21E-02 | 3,92E-02 |
| GOBP_REGULATION_OF_TRANSMEMBRANE_TRANSPORT | 479 | -1,369 | 4,21E-02 | 3,92E-02 |
| GOBP_ACUTE_PHASE_RESPONSE | 32 | -1,941 | 4,47E-02 | 4,17E-02 |
| GOBP_WNT_SIGNALING_PATHWAY_INVOLVED_IN_MIDBRAIN_DOPAMINERGIC_NEURON_DIFFERENTIATION | 12 | 1,926 | 4,47E-02 | 4,17E-02 |
| GOBP_RIBOSOMAL_SMALL_SUBUNIT_BIOGENESIS | 72 | -1,780 | 4,63E-02 | 4,32E-02 |
| GOBP_RELEASE_OF_SEQUESTERED_CALCIUM_ION_INTO_CYTOSOL_BY_ENDOPLASMIC_ RETICULUM | 29 | -1,978 | 4,79E-02 | 4,47E-02 |
| GOBP_RESPONSE_TO_INTERLEUKIN_1 | 111 | -1,699 | 4,79E-02 | 4,47E-02 |
| GOBP_REGULATION_OF_SIGNALING_RECEPTOR_ACTIVITY | 131 | -1,651 | 4,95E-02 | 4,62E-02 |

**Supplementary table 2: List of significant (p.adj < 0.05) differentially expressed genes (DEGs) between DMD organoids treated with µDystrophin (dEx8-9-µDys) and untreated (dEx8-9)**

| SYMBOL | baseMean | log2FoldChange | padj |
|---|---|---|---|
| LIG1 | 1000,5392 | 0,6531 | 5,83E-06 |
| MIR22HG | 1063,4607 | -0,6968 | 5,83E-06 |
| SOHLH2 | 576,2194 | -0,9356 | 8,75E-06 |
| NUP160 | 1050,8418 | 0,7835 | 3,34E-05 |
| PRKG1 | 1450,4331 | -0,7294 | 8,66E-05 |
| CAVIN1 | 5254,7199 | -0,6814 | 9,77E-05 |
| SLC37A4 | 626,1722 | 0,5484 | 1,15E-04 |
| ZMAT3 | 3171,2789 | 0,6578 | 1,56E-04 |
| OAS2 | 77,1247 | -1,9659 | 3,33E-04 |
| LIPG | 290,8780 | 1,0119 | 4,36E-04 |
| RHAG | 19,2703 | -2,8476 | 4,66E-04 |
| SELPLG | 193,8823 | -0,8892 | 4,66E-04 |
| PTCHD4 | 657,6944 | 0,7178 | 6,61E-04 |
| GLIPR2 | 659,5379 | -0,5602 | 6,93E-04 |
| KIFC3 | 1212,9565 | -0,5268 | 6,93E-04 |
| XRCC6 | 6453,6493 | 0,3962 | 6,93E-04 |
| ZNF197 | 559,6452 | 0,4867 | 6,93E-04 |
| GBP4 | 42,3899 | -1,5463 | 9,35E-04 |
| RBMX | 4139,7228 | 0,4207 | 9,35E-04 |
| NASP | 3361,1595 | 0,5295 | 9,69E-04 |
| POU3F4 | 67,5933 | 0,8698 | 9,83E-04 |
| POU3F3 | 126,0967 | 1,1402 | 1,03E-03 |
| CDC7 | 345,0627 | 0,7043 | 1,06E-03 |
| PLEKHD1 | 52,1518 | -1,0157 | 1,31E-03 |
| CCNF | 382,9842 | 0,5132 | 1,32E-03 |
| TGFB2-OT1 | 66,1781 | -8,1647 | 1,86E-03 |
| ZNF736 | 136,6268 | -1,9833 | 1,90E-03 |
| IFIT3 | 174,8642 | -2,0282 | 2,09E-03 |
| IRS3P | 69,3159 | -1,7759 | 2,23E-03 |
| IGF2BP1 | 4514,1792 | 0,6571 | 3,57E-03 |
| SPATA18 | 588,7327 | 1,1042 | 3,57E-03 |
| ZMYND8 | 1088,0392 | 0,5686 | 3,57E-03 |
| EDA2R | 725,5891 | 0,8696 | 3,73E-03 |
| CCDC69 | 161,8949 | -0,6429 | 3,73E-03 |
| NOS1 | 31,1763 | -2,9205 | 3,73E-03 |
| DYSF | 7420,6525 | -0,5755 | 4,05E-03 |
| ASS1 | 3601,5646 | -0,9537 | 4,31E-03 |
| POU5F1 | 109,6639 | -1,2288 | 4,65E-03 |
| FADS2 | 2164,0381 | 0,5955 | 4,82E-03 |
| EPHA7 | 690,0865 | 0,8938 | 5,18E-03 |
| GNRHR2 | 116,1791 | -1,4601 | 5,18E-03 |
| LOXL1-AS1 | 141,8355 | -1,2244 | 5,18E-03 |
| GOLGA7B | 48,5240 | -1,3446 | 6,54E-03 |
| C11orf96 | 180,9138 | -0,9547 | 6,97E-03 |
| CHAMP1 | 560,1796 | 0,5941 | 7,04E-03 |
| JUN-DT | 70,4570 | -1,2729 | 7,04E-03 |
| ATG4C | 134,2072 | 0,8595 | 7,24E-03 |
| CLN6 | 827,2692 | 0,4361 | 7,44E-03 |
| DGKB | 64,23627942 | -1,1852905 | 7,71E-03 |
| MPP4 | 27,2614601 | -1,6941284 | 7,84E-03 |
| OAS3 | 196,4213086 | -1,8024204 | 7,92E-03 |
| YPEL5 | 1201,664733 | -0,4021043 | 9,36E-03 |
| RRM2B | 2060,696768 | 0,6613807 | 9,62E-03 |
| LINC02897 | 76,44035369 | -1,3247572 | 1,02E-02 |
| CEBPD | 587,5727939 | -1,0087498 | 1,06E-02 |
| CDKN1A | 29145,72183 | 0,6142807 | 1,07E-02 |
| EPSTI1 | 126,7282082 | -1,3570121 | 1,07E-02 |
| MCM8 | 440,9408017 | 0,5645331 | 1,07E-02 |
| F13A1 | 1256,824034 | -1,4600273 | 1,11E-02 |
| NPPC | 28,13761912 | 1,259235 | 1,11E-02 |
| SKA2 | 550,1168461 | 0,4656071 | 1,11E-02 |
| LINC00943 | 29,83861433 | -1,4574996 | 1,12E-02 |
| MAP1LC3C | 295,4775752 | -0,7460999 | 1,12E-02 |
| PDIK1L | 134,48107 | 0,7817684 | 1,12E-02 |
| TBXA2R | 103,523345 | -1,0657371 | 1,12E-02 |
| WIPI1 | 2004,456144 | -0,4469893 | 1,12E-02 |
| PRR32 | 419,7438019 | -0,75543 | 1,13E-02 |
| SHFL | 338,9832493 | -1,111155 | 1,13E-02 |
| TLX3 | 51,04172393 | -1,0277524 | 1,13E-02 |
| TRIM24 | 1156,273009 | 0,5795963 | 1,13E-02 |
| UGP2 | 2436,639538 | -0,3658706 | 1,13E-02 |
| AIF1L | 905,0844211 | 0,4839651 | 1,15E-02 |
| CDC5L | 1134,788738 | 0,2518117 | 1,15E-02 |
| FBLIM1 | 1709,336299 | -0,5761398 | 1,15E-02 |
| FZD3 | 1130,016024 | 0,6218632 | 1,15E-02 |
| LINC-PINT | 139,4969872 | -1,0267185 | 1,15E-02 |
| MYADM | 3372,638902 | -0,4688729 | 1,15E-02 |
| MTTP | 45,88954342 | 0,7756902 | 1,26E-02 |
| CCNA2 | 292,8950383 | 0,6175241 | 1,28E-02 |
| FANCB | 35,22102936 | 1,0979167 | 1,35E-02 |
| SRPK1 | 828,0131426 | 0,6477474 | 1,35E-02 |
| HBA1 | 13,06940795 | -2,9004757 | 1,39E-02 |
| MIR503HG | 1765,733501 | -0,7430187 | 1,44E-02 |
| RN7SL525P | 17,87251773 | -1,5683208 | 1,46E-02 |

| SYMBOL | baseMean | log2FoldChang e | padj |
|---|---|---|---|
| RN7SL811P | 59,6374 | -1,5225 | 5,18E-03 |
| STON2 | 565,8415 | 0,4972 | 5,18E-03 |
| AOPEP | 459,9833 | -0,7111 | 5,31E-03 |
| CSRP3 | 2424,9814 | -1,0191 | 5,31E-03 |
| FRMPD1 | 522,2480 | -1,0549 | 5,31E-03 |
| MX1 | 152,4593 | -2,0200 | 5,31E-03 |
| PPIC | 926,5184 | -0,5455 | 5,31E-03 |
| USP44 | 129,2749 | 0,9939 | 5,31E-03 |
| ZNF454 | 42,4108 | -1,2562 | 5,31E-03 |
| ALDH1L1 | 82,8467 | -1,9848 | 5,33E-03 |
| ISG15 | 248,3680 | -1,5269 | 5,35E-03 |
| PCDHGA6 | 111,0031 | -1,0891 | 5,35E-03 |
| PPP1CC | 2943,4065 | 0,3492 | 5,35E-03 |
| NRIP3 | 168,0446 | -0,8902 | 5,36E-03 |
| ZNF37BP | 294,6567 | 0,7470 | 5,54E-03 |
| PCDHGC3 | 5251,3722 | -0,7528 | 5,89E-03 |
| INMT | 70,1100 | -1,6625 | 6,51E-03 |
| ZNF283 | 167,480938 | -1,41189398 | 1,87E-02 |
| ERMN | 27,0675747 | -1,57384389 | 1,90E-02 |
| LYNX1 | 112,282142 | 2,96788048 | 1,90E-02 |
| ODAD1 | 45,0083839 | -0,90639136 | 1,90E-02 |
| OR2A1-AS1 | 46,3106148 | -1,54670901 | 1,90E-02 |
| PSRC1 | 327,314093 | 0,60184032 | 1,90E-02 |
| BUB3 | 1547,54929 | 0,42872707 | 1,96E-02 |
| KIF21A | 709,972158 | 0,55028805 | 1,97E-02 |
| KLK10 | 25,9419403 | -1,5144194 | 1,97E-02 |
| RFC3 | 220,422361 | 0,56179904 | 1,97E-02 |
| ZCCHC24 | 2516,2161 | -0,39718062 | 1,98E-02 |
| DDX31 | 382,894563 | 0,38043155 | 2,01E-02 |
| HOXD8 | 116,94034 | -1,09757201 | 2,01E-02 |
| LIPC | 28,7927267 | -1,58787362 | 2,01E-02 |
| LIPH | 23,6618941 | -1,51538455 | 2,01E-02 |
| LINC00499 | 35,9908253 | -1,07008293 | 2,03E-02 |
| ARL6IP5 | 3119,8625 | -0,39684186 | 2,04E-02 |
| CTNNA2 | 164,344797 | 0,56661585 | 2,04E-02 |
| CSDE1 | 14212,3104 | 0,31028189 | 2,05E-02 |
| ILDR2 | 868,050967 | 1,08078504 | 2,05E-02 |
| LINC01091 | 31,4637231 | -1,34031157 | 2,05E-02 |
| MIR3945HG | 40,5285933 | -1,58092647 | 2,05E-02 |
| ATAD5 | 145,036264 | 0,55778651 | 2,06E-02 |
| KLC4 | 531,504412 | -0,49319453 | 2,07E-02 |
| FOXO6 | 101,783136 | -0,82350763 | 2,09E-02 |

| SYMBOL | baseMean | log2FoldChang e | padj |
|---|---|---|---|
| XRCC2 | 211,865771 | 0,57941254 | 1,46E-02 |
| IFIT1 | 175,35659 | -1,24446444 | 1,52E-02 |
| CIDECP1 | 12,2418102 | -21,4703528 | 1,60E-02 |
| PITX3 | 471,136095 | -0,62737975 | 1,61E-02 |
| LRRC43 | 20,8386894 | -1,24784732 | 1,63E-02 |
| PIP5K1B | 82,916257 | 0,68764793 | 1,65E-02 |
| CEP78 | 641,15656 | 0,53812082 | 1,68E-02 |
| TRIM7 | 228,962678 | -0,7467415 | 1,68E-02 |
| EDNRB | 296,254773 | 0,78888298 | 1,69E-02 |
| ESYT3 | 60,8344838 | -1,32147179 | 1,69E-02 |
| OAS1 | 253,934837 | -1,27461209 | 1,69E-02 |
| PYHIN1 | 24,0456204 | -1,45646901 | 1,71E-02 |
| SRSF10 | 2139,36676 | 0,36109586 | 1,71E-02 |
| UCA1-AS1 | 26,3118752 | -1,4921939 | 1,78E-02 |
| CPNE5 | 447,907943 | -0,67578407 | 1,79E-02 |
| POLE | 970,061897 | 0,74801259 | 1,79E-02 |
| SESN1 | 1215,68951 | 0,48138682 | 1,79E-02 |
| STING1 | 503,645733 | -0,7414322 | 2,56E-02 |
| ANP32A | 3389,51705 | 0,36538935 | 2,56E-02 |
| CDC6 | 261,235223 | 0,86998067 | 2,56E-02 |
| CDH4 | 235,957339 | 0,63232976 | 2,56E-02 |
| CERS6-AS1 | 209,0198 | -1,3998158 | 2,56E-02 |
| CMBL | 965,850471 | 0,45022932 | 2,56E-02 |
| HNRNPD | 5628,93922 | 0,36578071 | 2,56E-02 |
| NLRP10 | 16,1606333 | -1,60647097 | 2,56E-02 |
| SSRP1 | 3879,38071 | 0,25296334 | 2,56E-02 |
| TMEM176A | 24,1690012 | -1,5468932 | 2,56E-02 |
| PRXL2C | 345,925678 | -0,62565625 | 2,57E-02 |
| RARB | 46,947835 | -0,91977848 | 2,57E-02 |
| SOX18 | 28,9697216 | -1,68544517 | 2,57E-02 |
| BRI3BP | 264,834378 | 0,61473963 | 2,58E-02 |
| CCDC50 | 955,782014 | 0,36509167 | 2,58E-02 |
| CHI3L1 | 154,763257 | -1,01927984 | 2,58E-02 |
| GPR83 | 17,7632471 | -1,48932172 | 2,58E-02 |
| NUDT11 | 349,844894 | 0,36813909 | 2,58E-02 |
| SF3A3 | 2187,91601 | 0,38811093 | 2,58E-02 |
| SOX11 | 2998,83523 | 0,38974097 | 2,58E-02 |
| USP1 | 544,092255 | 0,46036528 | 2,58E-02 |
| PMP22 | 1018,85507 | -0,48992965 | 2,64E-02 |
| MYBPHL | 51,8185948 | -1,32327766 | 2,70E-02 |
| TMEM97 | 1182,11045 | 0,5197425 | 2,70E-02 |
| EEPD1 | 489,703333 | -0,55720839 | 2,70E-02 |

| SYMBOL | baseMean | log2FoldChange | padj |
|---|---|---|---|
| POLH | 1274,15352 | 0,56373593 | 2,09E-02 |
| GRM6 | 42,721619 | -0,89174221 | 2,10E-02 |
| ANKRD1 | 799,655446 | -0,84401649 | 2,13E-02 |
| FUT9 | 36,0394186 | 1,09392342 | 2,13E-02 |
| MTUS1 | 71,1237166 | 1,04469475 | 2,13E-02 |
| AQP6 | 13,7031049 | -1,40147273 | 2,17E-02 |
| EPB41 | 763,48974 | 0,52991359 | 2,17E-02 |
| FLACC1 | 15,4497271 | -1,55499984 | 2,17E-02 |
| LINC01291 | 18,2214416 | -2,75554674 | 2,17E-02 |
| MGP | 103,207147 | -1,70136348 | 2,17E-02 |
| QNG1 | 154,114192 | -0,86773983 | 2,17E-02 |
| FANCI | 599,901227 | 0,87069573 | 2,20E-02 |
| SPATA1 | 30,4436869 | -1,08899809 | 2,20E-02 |
| CXXC5 | 2991,76552 | -0,42588254 | 2,22E-02 |
| DHX58 | 37,664992 | -1,47538424 | 2,25E-02 |
| CBX5 | 3737,72288 | 0,46506731 | 2,26E-02 |
| HRK | 100,919519 | -0,85201675 | 2,27E-02 |
| LMNA | 17158,2938 | -0,41465111 | 2,27E-02 |
| NXPE3 | 540,530988 | 0,50815849 | 2,27E-02 |
| PDZRN3 | 1773,33403 | -0,4491143 | 2,27E-02 |
| STOX2 | 347,124414 | 0,72791639 | 2,27E-02 |
| METTL8 | 646,706111 | 0,64501018 | 2,31E-02 |
| RACGAP1 | 436,920539 | 0,73999513 | 2,31E-02 |
| TBC1D10A | 473,123966 | -0,42414148 | 2,31E-02 |
| ERCC6L | 66,2139124 | 0,70019767 | 2,32E-02 |
| SERBP1 | 8510,23578 | 0,24569264 | 2,32E-02 |
| DRAXIN | 740,673379 | 0,69388962 | 2,34E-02 |
| ODAD4 | 71,2214476 | 0,84997115 | 2,38E-02 |
| SHOX | 91,2163305 | -1,22492757 | 2,38E-02 |
| CHCHD2 | 1853,40946 | -0,82703332 | 2,41E-02 |
| SLC9A1 | 1105,79822 | -0,48602902 | 2,42E-02 |
| ADRA1A | 15,9256137 | -1,3991845 | 2,45E-02 |
| EZH2 | 806,462882 | 0,4990288 | 2,55E-02 |
| MAP3K7CL | 334,705811 | -0,73086497 | 2,55E-02 |
| SLC35F1 | 257,698992 | 0,51923616 | 2,55E-02 |
| TMPO | 1284,66103 | 0,62025084 | 2,55E-02 |
| USP18 | 53,3874799 | -1,2072564 | 2,74E-02 |
| WNK3 | 200,657631 | 0,66973085 | 2,76E-02 |
| PCDH19 | 235,086853 | 1,06829187 | 2,76E-02 |
| DSTN | 3204,74446 | -0,32961377 | 2,86E-02 |
| GMPS | 1381,26199 | 0,4438754 | 2,88E-02 |
| ACOT2 | 522,121924 | -0,82616567 | 2,88E-02 |

| SYMBOL | baseMean | log2FoldChange | padj |
|---|---|---|---|
| PELI1 | 252,217803 | 0,66295131 | 2,89E-02 |
| IFI44L | 78,2668217 | -2,10589865 | 2,96E-02 |
| KCTD1 | 382,201993 | 0,62304446 | 2,96E-02 |
| LINC00906 | 169,214111 | -1,41499517 | 2,96E-02 |
| P2RY6 | 145,023623 | -0,97115538 | 2,96E-02 |
| TRIM71 | 763,801797 | 0,74168532 | 2,96E-02 |
| MEX3A | 2635,24951 | 0,29546221 | 2,99E-02 |
| OR7A17 | 80,1726944 | -1,38364805 | 2,99E-02 |
| STEAP2 | 228,462796 | -1,06938105 | 2,99E-02 |
| TLCD2 | 414,25935 | -0,76385411 | 2,99E-02 |
| TGFBR1 | 1362,7939 | 0,58355195 | 3,03E-02 |
| ZNF676 | 47,0298453 | -1,40574889 | 3,27E-02 |
| LINC01284 | 43,4770399 | -1,26138533 | 3,33E-02 |
| TCF23 | 82,9552366 | -1,34111258 | 3,33E-02 |
| DBET | 10,5430892 | -2,04311204 | 3,36E-02 |
| DHX33 | 816,130284 | 0,57843117 | 3,36E-02 |
| NCAPG | 360,12047 | 0,74945051 | 3,36E-02 |
| PLK2 | 1649,53785 | 0,57936085 | 3,36E-02 |
| SALL2 | 1399,7216 | 0,52761439 | 3,36E-02 |
| SLITRK6 | 23,5975929 | 1,04935378 | 3,45E-02 |
| LGMN | 1188,1994 | -0,36114093 | 3,47E-02 |
| RN7SL693P | 9,10095369 | -1,80541088 | 3,48E-02 |
| KCNJ15 | 140,149492 | -1,28581717 | 3,49E-02 |
| HSPB6 | 551,668317 | -1,04794127 | 3,50E-02 |
| FUT2 | 47,6456569 | -1,3059679 | 3,55E-02 |
| MGLL | 1191,75611 | -0,54239522 | 3,55E-02 |
| KIF14 | 205,192034 | 0,74416251 | 3,57E-02 |
| MYT1L | 202,820634 | -0,9024693 | 3,57E-02 |
| KLHL23 | 489,315657 | 0,57867241 | 3,57E-02 |
| ACTL6A | 720,932895 | 0,49356944 | 3,58E-02 |
| CTSF | 184,30954 | -0,79632765 | 0,03579871 |
| LMCD1 | 602,458709 | -0,69652737 | 0,03579871 |
| NPNT | 1412,59973 | -0,66726875 | 0,03579871 |
| PA2G4 | 4159,10279 | 0,28081156 | 0,03579871 |
| TMEM74B | 93,9888354 | -0,6223597 | 0,03579871 |
| TRIM7-AS1 | 16,5424639 | -1,85560793 | 0,03579871 |
| IFIT2 | 64,9669999 | -1,84914502 | 0,03580412 |
| TRAC | 28,8670893 | -1,13382726 | 0,03596447 |
| MIOS | 416,698574 | 0,464336 | 0,03622571 |
| USH1C | 52,1065915 | -2,04301545 | 0,0363394 |
| ACTA1 | 16756,0636 | -0,7011716 | 0,03635759 |
| ZNF195 | 886,866103 | 0,48126523 | 0,03635759 |

| SYMBOL | baseMean | log2FoldChange | padj |
|---|---|---|---|
| PLP2 | 1317,47877 | -0,70731028 | 0,03683504 |
| PANTR1 | 27,8375799 | 1,41253013 | 3,72E-02 |
| USP3 | 715,6791 | 0,44532947 | 3,75E-02 |
| DNMT1 | 2786,88978 | 0,58887636 | 3,77E-02 |
| SH3BGRL2 | 188,445927 | 0,46191252 | 3,82E-02 |
| HMGB3 | 1505,11334 | 0,44760613 | 3,91E-02 |
| LINC01836 | 52,1355315 | -1,03278381 | 3,94E-02 |
| P3H2-AS1 | 32,4041892 | -1,19073158 | 3,97E-02 |
| ANGPT1 | 73,982223 | 0,77455461 | 4,02E-02 |
| ANXA2 | 10633,3481 | -0,59065215 | 4,02E-02 |
| TRPV3 | 118,117847 | 0,42670428 | 4,02E-02 |
| TSC22D2 | 943,745267 | 0,3835247 | 4,02E-02 |
| EXPH5 | 119,343359 | -0,84959224 | 4,03E-02 |
| HMGA2 | 885,551762 | 0,6920233 | 4,03E-02 |
| IPO9 | 2920,94477 | 0,4773307 | 4,03E-02 |
| RBL1 | 148,810088 | 0,58467359 | 4,03E-02 |
| RNF40 | 2671,4555 | 0,33783875 | 4,03E-02 |
| TNFRSF12A | 2365,62997 | -0,5009047 | 4,03E-02 |
| RAVER2 | 313,441009 | 0,5035075 | 4,05E-02 |
| TRPA1 | 14,8684788 | -3,69231915 | 4,05E-02 |
| ZNF561 | 1007,07414 | 0,51170599 | 4,05E-02 |
| DKC1 | 1213,93811 | 0,36284362 | 4,13E-02 |
| AGBL4 | 30,5218083 | -1,57919569 | 4,23E-02 |
| CA7 | 35,5226054 | -1,54103649 | 4,23E-02 |
| CDH8 | 272,014685 | 0,67907635 | 4,23E-02 |
| CYFIP2 | 2608,97205 | 0,40895033 | 4,23E-02 |
| GJC1 | 661,257461 | 0,55201593 | 4,23E-02 |
| MEIS1 | 349,994528 | 0,94374198 | 4,23E-02 |
| SLC30A3 | 101,989839 | -1,00801858 | 4,23E-02 |
| KRT80 | 75,5458366 | -0,8713149 | 4,23E-02 |
| TG | 35,7901253 | -2,09691612 | 4,32E-02 |
| SFPQ | 5378,37143 | 0,29981342 | 4,33E-02 |
| MCM7 | 2377,60618 | 0,47589273 | 4,34E-02 |
| POLD3 | 293,73598 | 0,57773064 | 4,34E-02 |
| CBS | 1440,41818 | 0,55047575 | 4,36E-02 |
| RN7SL274P | 23,1475721 | -1,26472716 | 4,36E-02 |
| SLC14A2 | 47,9952905 | -1,32891763 | 4,39E-02 |
| FBXO45 | 396,588789 | 0,33333201 | 4,40E-02 |

**Supplementary table 3: List of GO biological processes differentially expressed between DMD organoids treated with µDystrophin (dEx8-9-µDys) and isogenic control (IsoCTR)**

| Description | setSize | NES | p.adjust | qvalue |
|---|---|---|---|---|
| GOBP_CHROMOSOME_SEGREGATION | 315 | 2,441 | 2,438E-15 | 2,123E-15 |
| GOBP_DNA_CONFORMATION_CHANGE | 307 | 2,393 | 2,687E-14 | 2,340E-14 |
| GOBP_NUCLEAR_CHROMOSOME_SEGREGATION | 257 | 2,415 | 8,634E-14 | 7,519E-14 |
| GOBP_MITOTIC_NUCLEAR_DIVISION | 282 | 2,405 | 8,634E-14 | 7,519E-14 |
| GOBP_MUSCLE_CONTRACTION | 308 | -2,389 | 8,634E-14 | 7,519E-14 |
| GOBP_MUSCLE_SYSTEM_PROCESS | 390 | -2,271 | 9,797E-14 | 8,531E-14 |
| GOBP_SISTER_CHROMATID_SEGREGATION | 191 | 2,555 | 1,137E-13 | 9,904E-14 |
| GOBP_ORGANELLE_FISSION | 447 | 2,144 | 2,189E-13 | 1,907E-13 |
| GOBP_MITOTIC_SISTER_CHROMATID_SEGREGATION | 166 | 2,560 | 6,143E-13 | 5,349E-13 |
| GOBP_DNA_REPLICATION | 275 | 2,340 | 9,545E-13 | 8,312E-13 |
| GOBP_DNA_DEPENDENT_DNA_REPLICATION | 156 | 2,503 | 8,007E-12 | 6,972E-12 |
| GOBP_SPINDLE_ORGANIZATION | 199 | 2,294 | 8,586E-10 | 7,477E-10 |
| GOBP_DNA_REPLICATION_DEPENDENT_CHROMATIN_ORGANIZATION | 32 | 2,676 | 5,077E-09 | 4,421E-09 |
| GOBP_DNA_PACKAGING | 212 | 2,237 | 5,519E-09 | 4,806E-09 |
| GOBP_CELL_CYCLE_PHASE_TRANSITION | 487 | 1,883 | 6,046E-09 | 5,265E-09 |
| GOBP_RECOMBINATIONAL_REPAIR | 156 | 2,316 | 6,395E-09 | 5,569E-09 |
| GOBP_DOUBLE_STRAND_BREAK_REPAIR | 261 | 2,105 | 4,960E-08 | 4,319E-08 |
| GOBP_NEGATIVE_REGULATION_OF_MEGAKARYOCYTE_DIFFERENTIATION | 17 | 2,531 | 1,318E-07 | 1,147E-07 |
| GOBP_MITOTIC_CELL_CYCLE_PHASE_TRANSITION | 399 | 1,901 | 1,807E-07 | 1,573E-07 |
| GOBP_STRIATED_MUSCLE_CONTRACTION | 161 | -2,289 | 2,013E-07 | 1,753E-07 |
| GOBP_DNA_RECOMBINATION | 290 | 2,029 | 2,013E-07 | 1,753E-07 |
| GOBP_POSITIVE_REGULATION_OF_CELL_CYCLE | 285 | 1,993 | 2,667E-07 | 2,323E-07 |
| GOBP_SPINDLE_ASSEMBLY | 134 | 2,237 | 4,124E-07 | 3,591E-07 |
| GOBP_POSITIVE_REGULATION_OF_CELL_CYCLE_PROCESS | 211 | 2,086 | 4,124E-07 | 3,591E-07 |
| GOBP_REGULATION_OF_MITOTIC_CELL_CYCLE | 430 | 1,849 | 4,124E-07 | 3,591E-07 |
| GOBP_REGULATION_OF_SYSTEM_PROCESS | 482 | -1,866 | 5,351E-07 | 4,659E-07 |
| GOBP_AEROBIC_RESPIRATION | 166 | -2,210 | 6,655E-07 | 5,795E-07 |
| GOBP_DNA_REPLICATION_INDEPENDENT_CHROMATIN_ORGANIZATION | 34 | 2,490 | 7,647E-07 | 6,659E-07 |
| GOBP_MICROTUBULE_CYTOSKELETON_ORGANIZATION_INVOLVED_IN_MITOSIS | 150 | 2,167 | 9,443E-07 | 8,223E-07 |

| | | | | |
|---|---|---|---|---|
| GOBP_REGULATION_OF_MUSCLE_SYSTEM_PROCESS | 209 | -2,111 | 9,443E-07 | 8,223 E-07 |
| GOBP_OXIDATIVE_PHOSPHORYLATION | 119 | -2,294 | 1,544E-06 | 1,345 E-06 |
| GOBP_ENERGY_DERIVATION_BY_OXIDATION_OF_ORGANIC_C OMPOUNDS | 281 | -2,004 | 1,544E-06 | 1,345 E-06 |
| GOBP_CELLULAR_RESPIRATION | 208 | -2,124 | 2,099E-06 | 1,828 E-06 |
| GOBP_REGULATION_OF_MUSCLE_CONTRACTION | 143 | -2,244 | 2,512E-06 | 2,187 E-06 |
| GOBP_REGULATION_OF_CELL_CYCLE_PHASE_TRANSITION | 368 | 1,857 | 2,512E-06 | 2,187 E-06 |
| GOBP_ACTIVATION_OF_GTPASE_ACTIVITY | 107 | 2,221 | 3,770E-06 | 3,283 E-06 |
| GOBP_CHROMATIN_ASSEMBLY_OR_DISASSEMBLY | 200 | 2,012 | 3,873E-06 | 3,373 E-06 |
| GOBP_CYTOPLASMIC_TRANSLATION | 146 | -2,104 | 7,567E-06 | 6,590 E-06 |
| GOBP_PROTEIN_DNA_COMPLEX_ASSEMBLY | 172 | 2,044 | 7,567E-06 | 6,590 E-06 |
| GOBP_SKELETAL_MUSCLE_CONTRACTION | 41 | -2,400 | 9,857E-06 | 8,583 E-06 |
| GOBP_ATP_SYNTHESIS_COUPLED_ELECTRON_TRANSPORT | 77 | -2,246 | 1,091E-05 | 9,502 E-06 |
| GOBP_PROTEIN_DNA_COMPLEX_SUBUNIT_ORGANIZATION | 220 | 1,930 | 1,381E-05 | 1,203 E-05 |
| GOBP_DNA_GEOMETRIC_CHANGE | 94 | 2,232 | 1,515E-05 | 1,319 E-05 |
| GOBP_REGULATION_OF_MITOTIC_CELL_CYCLE_PHASE_TRANS ITION | 284 | 1,866 | 1,636E-05 | 1,425 E-05 |
| GOBP_SISTER_CHROMATID_COHESION | 53 | 2,312 | 2,002E-05 | 1,743 E-05 |
| GOBP_HEART_PROCESS | 224 | -1,990 | 2,002E-05 | 1,743 E-05 |
| GOBP_ACTIN_MYOSIN_FILAMENT_SLIDING | 14 | -2,278 | 2,131E-05 | 1,856 E-05 |
| GOBP_MITOCHONDRIAL_RESPIRATORY_CHAIN_COMPLEX_ASS EMBLY | 89 | -2,290 | 2,225E-05 | 1,937 E-05 |
| GOBP_CHROMOSOME_SEPARATION | 89 | 2,163 | 2,294E-05 | 1,998 E-05 |
| GOBP_REGULATION_OF_BLOOD_CIRCULATION | 219 | -2,011 | 2,350E-05 | 2,046 E-05 |
| GOBP_MICROTUBULE_ORGANIZING_CENTER_ORGANIZATION | 141 | 2,061 | 2,536E-05 | 2,208 E-05 |
| GOBP_TELOMERE_ORGANIZATION | 160 | 2,014 | 2,931E-05 | 2,552 E-05 |
| GOBP_REGULATION_OF_DNA_REPLICATION | 128 | 2,049 | 3,085E-05 | 2,687 E-05 |
| GOBP_MULTICELLULAR_ORGANISMAL_MOVEMENT | 53 | -2,319 | 3,613E-05 | 3,146 E-05 |
| GOBP_REGULATION_OF_CHROMOSOME_ORGANIZATION | 207 | 1,904 | 3,613E-05 | 3,146 E-05 |
| GOBP_HISTONE_MODIFICATION | 470 | 1,656 | 3,613E-05 | 3,146 E-05 |
| GOBP_CENTROMERE_COMPLEX_ASSEMBLY | 31 | 2,322 | 3,801E-05 | 3,310 E-05 |
| GOBP_REGULATION_OF_STRIATED_MUSCLE_CONTRACTION | 83 | -2,182 | 3,801E-05 | 3,310 E-05 |
| GOBP_CILIUM_ORGANIZATION | 384 | 1,733 | 3,801E-05 | 3,310 E-05 |
| GOBP_REGULATION_OF_CHROMOSOME_SEGREGATION | 82 | 2,195 | 3,914E-05 | 3,409 E-05 |
| GOBP_REGULATION_OF_HEART_CONTRACTION | 181 | -2,014 | 4,084E-05 | 3,556 E-05 |

| | | | | |
|---|---|---|---|---|
| GOBP_CELL_CYCLE_CHECKPOINT_SIGNALING | 165 | 1,968 | 4,886E-05 | 4,255 E-05 |
| GOBP_MITOTIC_CELL_CYCLE_CHECKPOINT_SIGNALING | 127 | 2,036 | 5,866E-05 | 5,108 E-05 |
| GOBP_ATP_METABOLIC_PROCESS | 240 | -1,881 | 6,521E-05 | 5,678 E-05 |
| GOBP_NADH_DEHYDROGENASE_COMPLEX_ASSEMBLY | 53 | -2,272 | 8,750E-05 | 7,619 E-05 |
| GOBP_NEURAL_PRECURSOR_CELL_PROLIFERATION | 136 | 1,997 | 9,374E-05 | 8,163 E-05 |
| GOBP_MEIOTIC_CELL_CYCLE | 215 | 1,842 | 9,621E-05 | 8,378 E-05 |
| GOBP_MEIOTIC_CELL_CYCLE_PROCESS | 164 | 1,912 | 1,334E-04 | 1,162 E-04 |
| GOBP_MITOTIC_SPINDLE_ORGANIZATION | 124 | 2,015 | 1,437E-04 | 1,251 E-04 |
| GOBP_PROTEIN_LOCALIZATION_TO_CHROMOSOME | 90 | 2,068 | 1,766E-04 | 1,538 E-04 |
| GOBP_REGULATION_OF_MEGAKARYOCYTE_DIFFERENTIATION | 35 | 2,251 | 1,879E-04 | 1,637 E-04 |
| GOBP_NEGATIVE_REGULATION_OF_CELL_CYCLE_PHASE_TRANSITION | 224 | 1,809 | 1,879E-04 | 1,637 E-04 |
| GOBP_CELL_CYCLE_DNA_REPLICATION | 43 | 2,218 | 1,948E-04 | 1,697 E-04 |
| GOBP_NEGATIVE_REGULATION_OF_CELL_CYCLE | 339 | 1,684 | 2,000E-04 | 1,742 E-04 |
| GOBP_MUSCLE_CELL_DEVELOPMENT | 168 | -1,916 | 2,240E-04 | 1,951 E-04 |
| GOBP_REGULATION_OF_CATION_TRANSMEMBRANE_TRANSPORT | 298 | -1,794 | 2,263E-04 | 1,970 E-04 |
| GOBP_RESPIRATORY_ELECTRON_TRANSPORT_CHAIN | 98 | -2,138 | 2,353E-04 | 2,049 E-04 |
| GOBP_CHROMATIN_REMODELING | 234 | 1,810 | 2,353E-04 | 2,049 E-04 |
| GOBP_STRIATED_MUSCLE_CELL_DEVELOPMENT | 64 | -2,244 | 2,922E-04 | 2,544 E-04 |
| GOBP_CELL_CELL_ADHESION_VIA_PLASMA_MEMBRANE_ADHESION_MOLECULES | 246 | 1,743 | 2,922E-04 | 2,544 E-04 |
| GOBP_CELLULAR_RESPONSE_TO_INTERLEUKIN_1 | 86 | -2,035 | 3,145E-04 | 2,738 E-04 |
| GOBP_MITOCHONDRIAL_ELECTRON_TRANSPORT_NADH_TO_UBIQUINONE | 43 | -2,231 | 3,763E-04 | 3,276 E-04 |
| GOBP_REGULATION_OF_DNA_DEPENDENT_DNA_REPLICATION | 51 | 2,164 | 4,143E-04 | 3,608 E-04 |
| GOBP_REGULATION_OF_CHROMOSOME_SEPARATION | 67 | 2,100 | 4,186E-04 | 3,645 E-04 |
| GOBP_NEGATIVE_REGULATION_OF_MITOTIC_CELL_CYCLE | 205 | 1,826 | 4,665E-04 | 4,062 E-04 |
| GOBP_REGULATION_OF_DENDRITIC_CELL_DIFFERENTIATION | 11 | -2,085 | 4,756E-04 | 4,142 E-04 |
| GOBP_NEGATIVE_REGULATION_OF_CELL_CYCLE_PROCESS | 266 | 1,719 | 4,988E-04 | 4,343 E-04 |
| GOBP_REGULATION_OF_DNA_METABOLIC_PROCESS | 376 | 1,600 | 5,115E-04 | 4,454 E-04 |
| GOBP_REGULATION_OF_MITOTIC_NUCLEAR_DIVISION | 104 | 1,953 | 5,195E-04 | 4,524 E-04 |
| GOBP_REGULATION_OF_NUCLEAR_DIVISION | 124 | 1,938 | 5,478E-04 | 4,770 E-04 |
| GOBP_NEUROMUSCULAR_PROCESS | 138 | -1,906 | 6,132E-04 | 5,340 E-04 |
| GOBP_NUCLEUS_ORGANIZATION | 129 | 1,886 | 6,132E-04 | 5,340 E-04 |
| GOBP_POSITIVE_REGULATION_OF_CELL_CYCLE_PHASE_TRANSITION | 109 | 1,926 | 6,133E-04 | 5,341 E-04 |

| | | | | |
|---|---|---|---|---|
| GOBP_GENERATION_OF_PRECURSOR_METABOLITES_AND_ENERGY | 436 | -1,642 | 6,135E-04 | 5,342E-04 |
| GOBP_NEGATIVE_REGULATION_OF_CHROMOSOME_ORGANIZATION | 86 | 1,985 | 6,897E-04 | 6,006E-04 |
| GOBP_SARCOMERE_ORGANIZATION | 40 | -2,200 | 7,359E-04 | 6,408E-04 |
| GOBP_HOMOPHILIC_CELL_ADHESION_VIA_PLASMA_MEMBRANE_ADHESION_MOLECULES | 157 | 1,870 | 7,359E-04 | 6,408E-04 |
| GOBP_DIVALENT_INORGANIC_CATION_HOMEOSTASIS | 401 | -1,656 | 7,406E-04 | 6,449E-04 |
| GOBP_PROTON_TRANSMEMBRANE_TRANSPORT | 123 | -2,009 | 8,395E-04 | 7,311E-04 |
| GOBP_SEX_DIFFERENTIATION | 234 | 1,735 | 8,522E-04 | 7,421E-04 |
| GOBP_DNA_REPLICATION_INITIATION | 36 | 2,150 | 8,635E-04 | 7,519E-04 |
| GOBP_CELL_CYCLE_G2_M_PHASE_TRANSITION | 150 | 1,819 | 8,660E-04 | 7,541E-04 |
| GOBP_DNA_INTEGRITY_CHECKPOINT_SIGNALING | 119 | 1,899 | 8,722E-04 | 7,595E-04 |
| GOBP_STRIATED_MUSCLE_CELL_DIFFERENTIATION | 251 | -1,745 | 8,722E-04 | 7,595E-04 |
| GOBP_ACTIN_MEDIATED_CELL_CONTRACTION | 94 | -1,963 | 8,894E-04 | 7,745E-04 |
| GOBP_CENTROSOME_DUPLICATION | 72 | 1,964 | 1,016E-03 | 8,847E-04 |
| GOBP_REGULATION_OF_DNA_REPAIR | 149 | 1,848 | 1,035E-03 | 9,012E-04 |
| GOBP_HETEROCHROMATIN_ORGANIZATION | 75 | 2,009 | 1,069E-03 | 9,305E-04 |
| GOBP_METAPHASE_ANAPHASE_TRANSITION_OF_CELL_CYCLE | 62 | 2,083 | 1,082E-03 | 9,420E-04 |
| GOBP_DNA_STRAND_ELONGATION_INVOLVED_IN_DNA_REPLICATION | 15 | 2,128 | 1,174E-03 | 1,022E-03 |
| GOBP_MITOTIC_SPINDLE_ASSEMBLY | 70 | 1,988 | 1,215E-03 | 1,058E-03 |
| GOBP_REGULATION_OF_CELL_CYCLE_G2_M_PHASE_TRANSITION | 104 | 1,890 | 1,368E-03 | 1,191E-03 |
| GOBP_REGULATION_OF_ION_TRANSMEMBRANE_TRANSPORT | 404 | -1,653 | 1,485E-03 | 1,293E-03 |
| GOBP_SARCOPLASMIC_RETICULUM_CALCIUM_ION_TRANSPORT | 36 | -2,140 | 1,553E-03 | 1,353E-03 |
| GOBP_NUCLEOSOME_ORGANIZATION | 146 | 1,828 | 1,599E-03 | 1,392E-03 |
| GOBP_DNA_DEPENDENT_DNA_REPLICATION_MAINTENANCE_OF_FIDELITY | 47 | 2,053 | 1,612E-03 | 1,404E-03 |
| GOBP_BLASTOCYST_GROWTH | 20 | 2,162 | 1,646E-03 | 1,434E-03 |
| GOBP_CARDIAC_MUSCLE_CONTRACTION | 123 | -1,968 | 1,650E-03 | 1,437E-03 |
| GOBP_NEGATIVE_REGULATION_OF_MITOTIC_CELL_CYCLE_PHASE_TRANSITION | 160 | 1,796 | 1,653E-03 | 1,439E-03 |
| GOBP_RESPONSE_TO_BACTERIUM | 454 | -1,607 | 1,653E-03 | 1,439E-03 |
| GOBP_MUSCLE_ORGAN_DEVELOPMENT | 314 | -1,680 | 1,723E-03 | 1,500E-03 |
| GOBP_MALE_SEX_DIFFERENTIATION | 140 | 1,796 | 1,725E-03 | 1,502E-03 |
| GOBP_HISTONE_METHYLATION | 148 | 1,793 | 1,725E-03 | 1,502E-03 |
| GOBP_POSITIVE_REGULATION_OF_MITOTIC_CELL_CYCLE | 114 | 1,889 | 1,730E-03 | 1,507E-03 |
| GOBP_NUCLEOSOME_ASSEMBLY | 99 | 1,916 | 1,804E-03 | 1,571E-03 |

| | | | | |
|---|---|---|---|---|
| GOBP_REGULATION_OF_METAL_ION_TRANSPORT | 340 | -1,676 | 1,842E-03 | 1,604 E-03 |
| GOBP_KINETOCHORE_ORGANIZATION | 23 | 2,143 | 1,856E-03 | 1,617 E-03 |
| GOBP_REGULATION_OF_DOUBLE_STRAND_BREAK_REPAIR_VIA_HOMOLOGOUS_RECOMBINATION | 68 | 1,977 | 1,878E-03 | 1,636 E-03 |
| GOBP_POSITIVE_REGULATION_OF_MITOTIC_CELL_CYCLE_PHASE_TRANSITION | 87 | 1,908 | 1,878E-03 | 1,636 E-03 |
| GOBP_INNER_CELL_MASS_CELL_PROLIFERATION | 13 | 2,079 | 1,965E-03 | 1,711 E-03 |
| GOBP_RESPONSE_TO_INTERLEUKIN_1 | 111 | -2,009 | 1,986E-03 | 1,729 E-03 |
| GOBP_DNA_UNWINDING_INVOLVED_IN_DNA_REPLICATION | 22 | 2,129 | 2,011E-03 | 1,751 E-03 |
| GOBP_CHROMOSOME_CONDENSATION | 37 | 2,098 | 2,508E-03 | 2,184 E-03 |
| GOBP_REGULATION_OF_SKELETAL_MUSCLE_ADAPTATION | 12 | -2,025 | 2,508E-03 | 2,184 E-03 |
| GOBP_POSITIVE_REGULATION_OF_DNA_METABOLIC_PROCESS | 221 | 1,637 | 2,826E-03 | 2,461 E-03 |
| GOBP_METAPHASE_PLATE_CONGRESSION | 66 | 1,971 | 3,015E-03 | 2,626 E-03 |
| GOBP_NEGATIVE_REGULATION_OF_GENE_EXPRESSION_EPIGENETIC | 74 | 1,920 | 3,073E-03 | 2,676 E-03 |
| GOBP_REGULATION_OF_EMBRYONIC_DEVELOPMENT | 60 | 1,952 | 3,157E-03 | 2,749 E-03 |
| GOBP_CALCIUM_ION_TRANSPORT | 353 | -1,582 | 3,157E-03 | 2,749 E-03 |
| GOBP_EOSINOPHIL_CHEMOTAXIS | 14 | -2,051 | 3,237E-03 | 2,818 E-03 |
| GOBP_NEUROBLAST_PROLIFERATION | 57 | 1,965 | 3,555E-03 | 3,096 E-03 |
| GOBP_ELECTRON_TRANSPORT_CHAIN | 148 | -1,775 | 3,555E-03 | 3,096 E-03 |
| GOBP_REGULATION_OF_TRANSMEMBRANE_TRANSPORT | 479 | -1,522 | 3,555E-03 | 3,096 E-03 |
| GOBP_MUSCLE_CELL_DIFFERENTIATION | 345 | -1,584 | 3,754E-03 | 3,269 E-03 |
| GOBP_PROTEIN_LOCALIZATION_TO_CHROMOSOME_CENTROMERIC_REGION | 25 | 2,053 | 3,917E-03 | 3,411 E-03 |
| GOBP_REPLICATION_FORK_PROCESSING | 39 | 2,019 | 3,917E-03 | 3,411 E-03 |
| GOBP_CELL_FATE_COMMITMENT | 215 | 1,653 | 3,917E-03 | 3,411 E-03 |
| GOBP_MITOTIC_DNA_INTEGRITY_CHECKPOINT_SIGNALING | 82 | 1,900 | 4,047E-03 | 3,524 E-03 |
| GOBP_ACUTE_INFLAMMATORY_RESPONSE | 78 | -1,982 | 4,172E-03 | 3,633 E-03 |
| GOBP_MITOTIC_METAPHASE_PLATE_CONGRESSION | 53 | 1,944 | 5,156E-03 | 4,489 E-03 |
| GOBP_REGULATION_OF_MITOTIC_SISTER_CHROMATID_SEGREGATION | 43 | 1,966 | 5,376E-03 | 4,681 E-03 |
| GOBP_SIGNAL_TRANSDUCTION_IN_RESPONSE_TO_DNA_DAMAGE | 168 | 1,683 | 5,739E-03 | 4,998 E-03 |
| GOBP_FOREBRAIN_DEVELOPMENT | 344 | 1,538 | 5,739E-03 | 4,998 E-03 |
| GOBP_DOUBLE_STRAND_BREAK_REPAIR_VIA_BREAK_INDUCED_REPLICATION | 12 | 2,036 | 5,869E-03 | 5,111 E-03 |
| GOBP_REGULATION_OF_CALCIUM_ION_TRANSPORT | 211 | -1,694 | 6,003E-03 | 5,227 E-03 |
| GOBP_SPINDLE_MIDZONE_ASSEMBLY | 11 | 1,991 | 6,141E-03 | 5,347 E-03 |
| GOBP_MIDBRAIN_DEVELOPMENT | 84 | 1,795 | 6,198E-03 | 5,397 E-03 |

| | | | | |
|---|---|---|---|---|
| GOBP_NEGATIVE_REGULATION_OF_LEUKOCYTE_MIGRATION | 41 | -1,975 | 6,215E-03 | 5,412 E-03 |
| GOBP_REGULATION_OF_DNA_DIRECTED_DNA_POLYMERASE_ ACTIVITY | 13 | 2,001 | 6,343E-03 | 5,524 E-03 |
| GOBP_MEGAKARYOCYTE_DIFFERENTIATION | 55 | 1,918 | 6,796E-03 | 5,918 E-03 |
| GOBP_MUSCLE_ADAPTATION | 98 | -1,893 | 6,796E-03 | 5,918 E-03 |
| GOBP_REGULATION_OF_DNA_RECOMBINATION | 120 | 1,734 | 6,796E-03 | 5,918 E-03 |
| GOBP_SENSORY_ORGAN_MORPHOGENESIS | 231 | 1,604 | 6,796E-03 | 5,918 E-03 |
| GOBP_LEUKOCYTE_MIGRATION | 293 | -1,603 | 6,796E-03 | 5,918 E-03 |
| GOBP_REGULATION_OF_GTPASE_ACTIVITY | 336 | 1,513 | 6,796E-03 | 5,918 E-03 |
| GOBP_CELLULAR_COMPONENT_ASSEMBLY_INVOLVED_IN_MO RPHOGENESIS | 99 | -1,908 | 6,934E-03 | 6,038 E-03 |
| GOBP_ACUTE_PHASE_RESPONSE | 32 | -2,016 | 7,004E-03 | 6,099 E-03 |
| GOBP_ACTIN_FILAMENT_BASED_MOVEMENT | 121 | -1,793 | 7,004E-03 | 6,099 E-03 |
| GOBP_REGULATION_OF_NEURAL_PRECURSOR_CELL_PROLIFE RATION | 79 | 1,803 | 7,426E-03 | 6,467 E-03 |
| GOBP_REGULATION_OF_SKELETAL_MUSCLE_CONTRACTION | 16 | -2,014 | 7,509E-03 | 6,539 E-03 |
| GOBP_LYMPHOCYTE_MIGRATION | 89 | -1,912 | 7,509E-03 | 6,539 E-03 |
| GOBP_NEPHRIC_DUCT_MORPHOGENESIS | 11 | 1,963 | 7,666E-03 | 6,676 E-03 |
| GOBP_CIRCULATORY_SYSTEM_PROCESS | 500 | -1,486 | 7,837E-03 | 6,824 E-03 |
| GOBP_CELL_CELL_SIGNALING_BY_WNT | 416 | 1,453 | 7,945E-03 | 6,918 E-03 |
| GOBP_MICROTUBULE_BUNDLE_FORMATION | 109 | 1,707 | 8,193E-03 | 7,134 E-03 |
| GOBP_WNT_SIGNALING_PATHWAY_INVOLVED_IN_MIDBRAIN_ DOPAMINERGIC_NEURON_DIFFERENTIATION | 12 | 2,008 | 8,324E-03 | 7,248 E-03 |
| GOBP_EOSINOPHIL_MIGRATION | 17 | -2,024 | 9,110E-03 | 7,933 E-03 |
| GOBP_NEUTROPHIL_MIGRATION | 90 | -1,822 | 9,150E-03 | 7,968 E-03 |
| GOBP_INTERLEUKIN_1_MEDIATED_SIGNALING_PATHWAY | 26 | -1,976 | 9,216E-03 | 8,025 E-03 |
| GOBP_REGULATION_OF_TRANSPORTER_ACTIVITY | 246 | -1,618 | 9,652E-03 | 8,405 E-03 |
| GOBP_MITOTIC_SISTER_CHROMATID_COHESION | 26 | 2,015 | 1,022E-02 | 8,900 E-03 |
| GOBP_REGULATION_OF_HIPPO_SIGNALING | 21 | 1,998 | 1,022E-02 | 8,900 E-03 |
| GOBP_REGULATION_OF_GENE_EXPRESSION_EPIGENETIC | 130 | 1,713 | 1,048E-02 | 9,122 E-03 |
| GOBP_ATTACHMENT_OF_SPINDLE_MICROTUBULES_TO_KINET OCHORE | 37 | 1,961 | 1,089E-02 | 9,487 E-03 |
| GOBP_POSITIVE_REGULATION_OF_GTPASE_ACTIVITY | 250 | 1,557 | 1,107E-02 | 9,639 E-03 |
| GOBP_EMBRYONIC_PATTERN_SPECIFICATION | 56 | 1,849 | 1,115E-02 | 9,714 E-03 |
| GOBP_REGULATION_OF_ORGANELLE_ASSEMBLY | 220 | 1,580 | 1,152E-02 | 1,003 E-02 |
| GOBP_NEGATIVE_REGULATION_OF_METAPHASE_ANAPHASE_ TRANSITION_OF_CELL_CYCLE | 40 | 1,941 | 1,163E-02 | 1,012 E-02 |

| | | | | |
|---|---|---|---|---|
| GOBP_REGULATION_OF_SMOOTHENED_SIGNALING_PATHWAY | 78 | 1,829 | 1,163E-02 | 1,012 E-02 |
| GOBP_REPRODUCTIVE_SYSTEM_DEVELOPMENT | 374 | 1,450 | 1,163E-02 | 1,012 E-02 |
| GOBP_DNA_REPLICATION_CHECKPOINT_SIGNALING | 16 | 1,999 | 1,177E-02 | 1,025 E-02 |
| GOBP_REGULATION_OF_CALCIUM_ION_TRANSMEMBRANE_TRANSPORT | 131 | -1,738 | 1,177E-02 | 1,025 E-02 |
| GOBP_PROTEIN_AUTOPHOSPHORYLATION | 214 | 1,576 | 1,201E-02 | 1,046 E-02 |
| GOBP_POSITIVE_REGULATION_OF_CYTOKINE_PRODUCTION | 373 | -1,521 | 1,201E-02 | 1,046 E-02 |
| GOBP_DNA_STRAND_ELONGATION | 37 | 1,949 | 1,216E-02 | 1,059 E-02 |
| GOBP_UROGENITAL_SYSTEM_DEVELOPMENT | 327 | 1,464 | 1,216E-02 | 1,059 E-02 |
| GOBP_REGULATION_OF_SISTER_CHROMATID_COHESION | 21 | 1,963 | 1,330E-02 | 1,158 E-02 |
| GOBP_PROTEIN_LOCALIZATION_TO_CHROMATIN | 30 | 1,958 | 1,330E-02 | 1,158 E-02 |
| GOBP_REGULATION_OF_ATPASE_COUPLED_CALCIUM_TRANSMEMBRANE_TRANSPORTER_ACTIVITY | 12 | -1,905 | 1,330E-02 | 1,158 E-02 |
| GOBP_SPINAL_CORD_DEVELOPMENT | 87 | 1,737 | 1,330E-02 | 1,158 E-02 |
| GOBP_OSSIFICATION | 371 | 1,425 | 1,330E-02 | 1,158 E-02 |
| GOBP_REGULATION_OF_CHROMOSOME_CONDENSATION | 11 | 1,929 | 1,335E-02 | 1,163 E-02 |
| GOBP_REGULATION_OF_ATP_DEPENDENT_ACTIVITY | 70 | -1,794 | 1,336E-02 | 1,164 E-02 |
| GOBP_CARDIAC_MUSCLE_TISSUE_MORPHOGENESIS | 55 | -1,866 | 1,349E-02 | 1,175 E-02 |
| GOBP_REGULATION_OF_CALCIUM_ION_TRANSMEMBRANE_TRANSPORTER_ACTIVITY | 81 | -1,801 | 1,349E-02 | 1,175 E-02 |
| GOBP_CHROMOSOME_LOCALIZATION | 80 | 1,782 | 1,408E-02 | 1,226 E-02 |
| GOBP_MEIOTIC_CHROMOSOME_SEGREGATION | 74 | 1,778 | 1,421E-02 | 1,237 E-02 |
| GOBP_DNA_TEMPLATED_TRANSCRIPTION_INITIATION | 126 | 1,636 | 1,465E-02 | 1,276 E-02 |
| GOBP_INTERLEUKIN_6_PRODUCTION | 125 | -1,731 | 1,468E-02 | 1,278 E-02 |
| GOBP_NEGATIVE_REGULATION_OF_INNATE_IMMUNE_RESPONSE | 62 | -1,890 | 1,519E-02 | 1,323 E-02 |
| GOBP_CYTOCHROME_COMPLEX_ASSEMBLY | 35 | -1,943 | 1,567E-02 | 1,365 E-02 |
| GOBP_RESPONSE_TO_INORGANIC_SUBSTANCE | 470 | -1,453 | 1,581E-02 | 1,377 E-02 |
| GOBP_CHROMOSOME_ORGANIZATION_INVOLVED_IN_MEIOTIC_CELL_CYCLE | 54 | 1,825 | 1,681E-02 | 1,464 E-02 |
| GOBP_DEVELOPMENT_OF_PRIMARY_SEXUAL_CHARACTERISTICS | 192 | 1,588 | 1,681E-02 | 1,464 E-02 |
| GOBP_HIPPO_SIGNALING | 40 | 1,899 | 1,711E-02 | 1,490 E-02 |
| GOBP_STRIATED_MUSCLE_ADAPTATION | 42 | -1,961 | 1,762E-02 | 1,535 E-02 |
| GOBP_NEGATIVE_REGULATION_OF_NUCLEAR_DIVISION | 52 | 1,883 | 1,772E-02 | 1,543 E-02 |
| GOBP_MAINTENANCE_OF_CELL_NUMBER | 132 | 1,671 | 1,901E-02 | 1,655 E-02 |
| GOBP_CELL_DIFFERENTIATION_IN_SPINAL_CORD | 41 | 1,882 | 1,915E-02 | 1,668 E-02 |
| GOBP_MEIOSIS_I_CELL_CYCLE_PROCESS | 106 | 1,703 | 1,936E-02 | 1,686 E-02 |

| | | | | |
|---|---|---|---|---|
| GOBP_MORPHOGENESIS_OF_AN_EPITHELIAL_FOLD | 20 | 1,961 | 1,974E-02 | 1,719 E-02 |
| GOBP_INTERSTRAND_CROSS_LINK_REPAIR | 31 | 1,848 | 2,026E-02 | 1,764 E-02 |
| GOBP_ATP_BIOSYNTHETIC_PROCESS | 48 | -1,885 | 2,028E-02 | 1,766 E-02 |
| GOBP_MITOTIC_G2_M_TRANSITION_CHECKPOINT | 47 | 1,824 | 2,028E-02 | 1,766 E-02 |
| GOBP_MONONUCLEAR_CELL_MIGRATION | 148 | -1,640 | 2,132E-02 | 1,857 E-02 |
| GOBP_CALCIUM_ION_TRANSMEMBRANE_TRANSPORT | 264 | -1,530 | 2,180E-02 | 1,898 E-02 |
| GOBP_LYMPHOCYTE_CHEMOTAXIS | 40 | -1,874 | 2,197E-02 | 1,913 E-02 |
| GOBP_SKELETAL_MUSCLE_ADAPTATION | 21 | -1,927 | 2,246E-02 | 1,955 E-02 |
| GOBP_POSITIVE_REGULATION_OF_TRANSFERASE_ACTIVITY | 499 | 1,359 | 2,246E-02 | 1,955 E-02 |
| GOBP_CATECHOLAMINE_SECRETION | 47 | -1,860 | 2,255E-02 | 1,963 E-02 |
| GOBP_RESPONSE_TO_STIMULUS_INVOLVED_IN_REGULATION_OF_MUSCLE_ADAPTATION | 13 | -1,915 | 2,276E-02 | 1,982 E-02 |
| GOBP_PROTEIN_LOCALIZATION_TO_CONDENSED_CHROMOSOME | 19 | 2,013 | 2,297E-02 | 2,000 E-02 |
| GOBP_PATTERN_SPECIFICATION_PROCESS | 398 | 1,427 | 2,386E-02 | 2,078 E-02 |
| GOBP_CELL_CHEMOTAXIS | 233 | -1,573 | 2,440E-02 | 2,125 E-02 |
| GOBP_REGULATION_OF_ATTACHMENT_OF_SPINDLE_MICROTUBULES_TO_KINETOCHORE | 13 | 1,888 | 2,445E-02 | 2,129 E-02 |
| GOBP_POSITIVE_REGULATION_OF_INTERLEUKIN_1_PRODUCTION | 60 | -1,876 | 2,445E-02 | 2,129 E-02 |
| GOBP_POSITIVE_REGULATION_OF_NERVOUS_SYSTEM_DEVELOPMENT | 250 | 1,494 | 2,445E-02 | 2,129 E-02 |
| GOBP_FAT_SOLUBLE_VITAMIN_METABOLIC_PROCESS | 30 | 1,889 | 2,458E-02 | 2,140 E-02 |
| GOBP_HISTONE_H3_K4_METHYLATION | 67 | 1,756 | 2,458E-02 | 2,140 E-02 |
| GOBP_CELL_CYCLE_G1_S_PHASE_TRANSITION | 217 | 1,517 | 2,491E-02 | 2,170 E-02 |
| GOBP_POSITIVE_REGULATION_OF_HISTONE_METHYLATION | 45 | 1,845 | 2,555E-02 | 2,225 E-02 |
| GOBP_REGULATION_OF_CYTOSOLIC_CALCIUM_ION_CONCENTRATION | 264 | -1,517 | 2,556E-02 | 2,226 E-02 |
| GOBP_POSITIVE_REGULATION_OF_INTERLEUKIN_1_BETA_PRODUCTION | 51 | -1,841 | 2,556E-02 | 2,226 E-02 |
| GOBP_VENTRICULAR_CARDIAC_MUSCLE_TISSUE_MORPHOGENESIS | 44 | -1,864 | 2,562E-02 | 2,231 E-02 |
| GOBP_REGULATION_OF_DOUBLE_STRAND_BREAK_REPAIR | 107 | 1,683 | 2,575E-02 | 2,242 E-02 |
| GOBP_SKIN_DEVELOPMENT | 194 | 1,515 | 2,575E-02 | 2,242 E-02 |
| GOBP_CHROMATIN_REMODELING_AT_CENTROMERE | 11 | 1,886 | 2,653E-02 | 2,310 E-02 |
| GOBP_PEPTIDYL_LYSINE_METHYLATION | 134 | 1,650 | 2,685E-02 | 2,338 E-02 |
| GOBP_PROTEIN_METHYLATION | 185 | 1,511 | 2,685E-02 | 2,338 E-02 |
| GOBP_DEFENSE_RESPONSE_TO_BACTERIUM | 159 | -1,652 | 2,814E-02 | 2,451 E-02 |
| GOBP_RESPONSE_TO_PEPTIDOGLYCAN | 10 | -1,844 | 2,864E-02 | 2,494 E-02 |

| | | | | |
|---|---|---|---|---|
| GOBP_POSITIVE_REGULATION_OF_DOUBLE_STRAND_BREAK_REPAIR_VIA_HOMOLOGOUS_RECOMBINATION | 35 | 1,849 | 2,872E-02 | 2,501 E-02 |
| GOBP_VENTRICULAR_CARDIAC_MUSCLE_TISSUE_DEVELOPMENT | 52 | -1,882 | 2,893E-02 | 2,519 E-02 |
| GOBP_SENSORY_ORGAN_DEVELOPMENT | 490 | 1,340 | 2,904E-02 | 2,529 E-02 |
| GOBP_MITOTIC_CHROMOSOME_CONDENSATION | 19 | 1,983 | 2,933E-02 | 2,554 E-02 |
| GOBP_KINETOCHORE_ASSEMBLY | 18 | 1,933 | 3,074E-02 | 2,676 E-02 |
| GOBP_REGULATION_OF_LEUKOCYTE_MIGRATION | 171 | -1,593 | 3,096E-02 | 2,696 E-02 |
| GOBP_RESPONSE_TO_VITAMIN_A | 12 | 1,898 | 3,101E-02 | 2,701 E-02 |
| GOBP_NEPHRON_DEVELOPMENT | 137 | 1,540 | 3,130E-02 | 2,726 E-02 |
| GOBP_POSITIVE_REGULATION_OF_SMOOTHENED_SIGNALING_PATHWAY | 34 | 1,862 | 3,217E-02 | 2,801 E-02 |
| GOBP_NEGATIVE_REGULATION_OF_CALCIUM_ION_TRANSPORT | 48 | -1,813 | 3,251E-02 | 2,831 E-02 |
| GOBP_SMOOTHENED_SIGNALING_PATHWAY | 134 | 1,626 | 3,456E-02 | 3,009 E-02 |
| GOBP_REGULATION_OF_MEIOTIC_CELL_CYCLE | 40 | 1,805 | 3,470E-02 | 3,022 E-02 |
| GOBP_T_CELL_MIGRATION | 52 | -1,845 | 3,685E-02 | 3,209 E-02 |
| GOBP_REGULATION_OF_HISTONE_METHYLATION | 73 | 1,707 | 3,767E-02 | 3,281 E-02 |
| GOBP_STEM_CELL_PROLIFERATION | 59 | 1,720 | 3,958E-02 | 3,446 E-02 |
| GOBP_PROTEIN_ACETYLATION | 202 | 1,520 | 3,958E-02 | 3,446 E-02 |
| GOBP_LONG_CHAIN_FATTY_ACID_BIOSYNTHETIC_PROCESS | 22 | 1,885 | 4,020E-02 | 3,501 E-02 |
| GOBP_KIDNEY_EPITHELIUM_DEVELOPMENT | 132 | 1,611 | 4,023E-02 | 3,503 E-02 |
| GOBP_POSITIVE_REGULATION_OF_NEUROGENESIS | 206 | 1,521 | 4,033E-02 | 3,512 E-02 |
| GOBP_HOMOLOGOUS_RECOMBINATION | 49 | 1,709 | 4,087E-02 | 3,559 E-02 |
| GOBP_FEMALE_SEX_DIFFERENTIATION | 107 | 1,635 | 4,105E-02 | 3,574 E-02 |
| GOBP_SPINAL_CORD_ASSOCIATION_NEURON_DIFFERENTIATION | 10 | 1,836 | 4,246E-02 | 3,698 E-02 |
| GOBP_MORPHOGENESIS_OF_AN_EPITHELIAL_BUD | 15 | 1,843 | 4,461E-02 | 3,884 E-02 |
| GOBP_EPIDERMIS_MORPHOGENESIS | 29 | 1,794 | 4,859E-02 | 4,231 E-02 |
| GOBP_PEPTIDYL_LYSINE_ACETYLATION | 179 | 1,509 | 4,859E-02 | 4,231 E-02 |
| GOBP_REGULATION_OF_NEUROGENESIS | 334 | 1,395 | 4,859E-02 | 4,231 E-02 |

**Supplementary table 4: Cell lines list**

| | Current name | Reference | Healthy individuals and patients |
|---|---|---|---|
| **Human induced pluripotent stem cells (hiPSCs)** | CTR1 | AG08C5 Coriell | Skin fibroblast, 1-year-old healthy male |
| | CTR2 | M180 CECS | Skeletal myoblasts, healthy male |
| | IsoCTR | *Kyrychenko et al 2017* | Peripheral blood mononuclear cells healthy male |
| | DMDdEx45 | GM25313 Coriell | Skin fibroblast, 13-year-old DMD male patient, deletion ex45 |
| | DMDdEx8-9 | *Kyrychenko et al 2017* | Peripheral blood mononuclear cells healthy male |
| | DMDdEx8-43 | M202 | Skeletal myoblasts, DMD male patient, out-of-frame deletion exon 8-43 |
| **Immortalized skin fibroblasts (hTERT/CDK4)** | CTR fibroblasts | AB1191C16PV | Skin, 16-year-old healthy male |
| | DMD fibroblasts | AB1024DMD11Q | , Skin 11-year-old DMD male, stop-codon exon 59 |
| **Primary skin fibroblasts** | Fibroblasts control 1 | 5-8898 DNA Bank Genethon | Skin fibroblasts healthy patient |
| | Fibroblasts control 2 | 11-1610 DNA Bank Genethon | Skin fibroblasts healthy patient |
| | Fibroblasts control 3 | 11-1973 DNA Bank Genethon | Skin fibroblasts healthy patient |
| | Fibroblasts DMD 1 | 5-11618 DNA Bank Genethon | Skin fibroblasts DMD patient |
| | Fibroblasts DMD 2 | 5-12608 DNA Bank Genethon | Skin fibroblasts DMD patient |
| | Fibroblasts DMD 3 | 5-11613 DNA Bank Genethon | Skin fibroblasts DMD patient |

**Supplementary table 5: Antibodies used for immunofluorescence and capillary western blot**

| Antibodies | Supplier | Reference |
|---|---|---|
| Mouse monoclonal Myosin Heavy Chain (used at 1:10) | DSHB | Cat#MF20, RRID:AB_2147781 |
| Rabbit polyclonal Vimentin (used at 1:100) | Proteintech | Cat#10366-1-AP, RRID:AB_2273020 |
| Mouse monoclonal Dystrophin (used at 1:20) | Leica Biosystem | Cat#NCL-DYSB, RRID:AB_563691 |
| Mouse monoclonal Fibronectin (used at 1:200) | Sigma-Aldrich | Cat# F7387, RRID:AB_476988 |
| Rabbit monoclonal pSMAD-3 (used at 1:100) | Abcam | Cat# ab52903, RRID:AB_882596 |
| Mouse monoclonal α-Dystroglycan (used at 1:100) | Millipore | Cat# 05-298, RRID:AB_309674 |
| Mouse monoclonal β-Dystroglycan (used at 1:100) | Leica Biosystems | Cat#NCL-b-DG, RRID:AB_442043 |
| Rabbit monoclonal alpha-actinin 2 (used at 1:100) | Thermo Fisher Scientific | Cat# 701914, RRID:AB_2688290 |
| Mouse monoclonal Oct3/4 (used at 1:200) | Santa Cruz Biotechnology | Cat# sc-5279, RRID: AB_628051 |
| Mouse monoclonal Pax7 (used at 1:100) | DSHB | Cat# PAX7, RRID:AB_2299243 |
| Goat anti-Mouse IgG1 Cross-Adsorbed Secondary Antibody, Alexa Fluor 594 | Thermo Fisher Scientific | Cat#A-21125, RRID:AB_2535767 |
| Goat anti-Rabbit IgG (Heavy chain), Superclonal™ Recombinant Secondary Antibody, Alexa Fluor 488 | Thermo Fisher Scientific | Cat# A27034, RRID:AB_2536097 |
| Goat anti-Mouse IgG2a Cross-Adsorbed Secondary Antibody, Alexa Fluor 594 | Thermo Fisher Scientific | Cat#A-21135, RRID:AB_2535774 |

| | | |
|---|---|---|
| Goat anti-Mouse IgG2b Cross-Adsorbed Secondary Antibody, Alexa Fluor 647 | Thermo Fisher Scientific | Cat#A-21242, RRID:AB_2535811 |

**Supplementary table 6: List of primers used for viral copy number analysis and gene expression**

| Primers for gene expression analysis | | |
|---|---|---|
| **Gene** | **Forward** | **Reverse** |
| GAPDH | ACAACTTTGGTATCGTGGAAGG | GCCATCACGCCACAGTTTC |
| MYOGENIN | AATGCAGCTCTCACAGCGCCTC | TCAGCCGTGAGCAGATGATCC |
| MCK | TGGAGAAGCTCTCTGTGGAAGCT | TCCGTCATGCTCTTCAGAGGGTA |
| MYH2 | AGAAACTTCGCATGGACCTAGA | CCAAGTGCCTGTTCATCTTCA |
| MYH7 | ACTGCCGAGACCGAGTATG | GCGATCCTTGAGGTTGTAGAGC |
| μDYS | GCACCACCAGATGCACTAT | GTGTAGGCGTAGCTCTTGAAT |
| COL1A1 | AAGAGGAAGGCCAAGTCGAG | GTTTCCACACGTCTCGGTCA |
| FN1 | AGCCGAGGTTTTAACTGCGA | CCCACTCGGTAAGTGTTCCC |
| TGF-β1 | GCCTGAGGCCGACTACTA | CTGTGTGTACTCTGCTTGAACT |
| **Primers for VCN analysis** | | |
| **Gene** | **Forward** | **Reverse** |
| P0 | CTCCAAGCAGATGCAGCAGA | ATAGCCTTGCGCATCATGGT |
| μDYS | GCACCACCAGATGCACTAT | GTGTAGGCGTAGCTCTTGAAT |

**Supplementary table 7: ELISA kit used for secreted protein analysis**

| **Protein target** | **Supplier** | **Reference** |
|---|---|---|
| Fibronectin 1 | Abcam | ab219046 |
| TGF-β1 | Abcam | ab100647 |
| Collagen IV | Bio-techne | NBP2-75864 |
